# Sex-biased expression is associated with chromatin state in *D. melanogaster* and *D. simulans*

**DOI:** 10.1101/2023.01.13.523946

**Authors:** Adalena V. Nanni, Natalie Martinez, Rita Graze, Alison Morse, Jeremy R. B. Newman, Vaibhav Jain, Srna Vlaho, Sarah Signor, Sergey V. Nuzhdin, Rolf Renne, Lauren M. McIntyre

## Abstract

We propose a new model for the association of chromatin state and sex-bias in expression. We hypothesize enrichment of open chromatin in the sex where we see expression bias (OS) and closed chromatin in the opposite sex (CO). In this study of *D. melanogaster* and *D. simulans* head tissue, sex-bias in expression is associated with H3K4me3 (open mark) in males for male-biased genes and in females for female-biased genes in both species. Sex-bias in expression is also largely conserved in direction and magnitude between the two species on the X and autosomes. In male-biased orthologs, the sex-bias ratio is more divergent between species if both species have H3K27me2me3 marks in females compared to when either or neither species has H3K27me2me3 in females. H3K27me2me3 marks in females are associated with male-bias in expression on the autosomes in both species, but on the X only in *D. melanogaster*. In female-biased orthologs the relationship between the species for the sex-bias ratio is similar regardless of the H3K27me2me3 marks in males. Female-biased orthologs are more similar in the ratio of sex-bias than male-biased orthologs and there is an excess of male-bias in expression in orthologs that gain/lose sex-bias. There is an excess of male-bias in sex-limited expression in both species suggesting excess male-bias is due to rapid evolution between the species. The X chromosome has an enrichment in male-limited H3K4me3 in both species and an enrichment of sex-bias in expression compared to the autosomes.

## Introduction

Chromatin accessibility is known to be important for multiple levels of gene regulation, as well as in large scale modifications of expression such as in dosage compensation of sex chromosomes. H3K4me3 is an open chromatin mark correlated with activate expression (Santos-Rosa, et al. 2002; Schneider, et al. 2004) and closed chromatin marks H3K27me2 and H3K27me3, together referred to as H3K27me2me3, are correlated with silenced expression (Wang, et al. 2008; Juan, et al. 2016). These three marks act together with other histone modifications, DNA methylation, and chromatin factors, in the establishment and modification of chromatin accessibility (reviewed in Boros 2012). and the extent to which epigenetics influences behavior is an emerging paradigm explored in several systems (e.g., Spannhoff, et al. 2011; Sun, et al. 2015; Elliott, et al. 2016; Sun, et al. 2016; Opachaloemphan, et al. 2018; Qin, et al. 2018; Bludau, et al. 2019) including in plastic behavioral traits such as foraging (Anreiter, et al. 2017; Anreiter and Sokolowski 2019).

Differences between the X chromosome and autosomes in the evolution of gene expression may be due to changes in regulation of chromatin conformation. There is evidence in third instar larvae for an enrichment of open chromatin marks on the X chromosome compared to the autosomes in both males and females of *D. miranda* and *D. melanogaster*, as well as more open chromatin in males compared to females on the X chromosome and more closed chromatin in females compared to males (Brown and Bachtrog 2014). Furthermore, Brown, et al. (Brown, et al. 2020) also demonstrated that the male Y chromosome has a genome-wide effect on heterochromatin factors, leading to a heterochromatin sink effect. Specifically, the male X chromosome and autosomes are more open compared to the female due to a sequestering of closed chromatin factors on the Y chromosome (Henikoff 1996; Francisco and Lemos 2014).

Chromatin remodeling genes have known roles in sex determination and may be important for overall regulation of sex differences in gene expression. For example, the sex determination gene *fru* aids in recruiting of histone deacetylase (HDAC) *Rpd3* and heterochromatin protein 1A (HP1a) encoded by *Su(var)205* (Ito, et al. 2012) to genes associated with male courtship behaviors. The expression of *fru* decreases with mutation of a histone demethylase *kdm4* (Lorbeck, et al. 2010), resulting in the male chain mating phenotype, also found in *fru* mutants (Ito, et al. 1996). Sexually dimorphic chromatin modifications such as H3K9me2 (associated with closed chromatin) and H4K16ac (associated with open chromatin) have been reported (Brown and Bachtrog 2014).

In *D. melanogaster*, sexually dimorphic chromatin accessibility is stage and cell-type specific (Palmateer, et al. 2021). For example, in *fru-P1-expressing* neurons of 1-day old adults the TSS of genes are enriched for H3K4me3 in males compared to females, while the reverse is true in 10- to 12-day old adults. These sex differences at the TSS were not observed in *elav*-expressing neurons (Palmateer, et al. 2021) supporting their role in directing sex-specific behaviors (Ito, et al. 1996; Ryner, et al. 1996; Demir and Dickson 2005; Manoli, et al. 2005; Stockinger, et al. 2005; Goldman and Arbeitman 2007; reviewed in Yamamoto and Koganezawa 2013).

Morphological and behavioral differences between males and females are common in sexually reproducing organisms (Hedrick and Temeles 1989). *Drosophila* species have diverged in sexually dimorphic morphology and behaviors (Sturtevant 1920; Ewing and Bennet-Clark 1968; Cobb, et al. 1989; Chakir, et al. 2002; Kopp 2011; Arthur, et al. 2013). For example, there has been relatively rapid diversification in reproductive behaviors such as the male courtship song (e.g., Ritchie, et al. 1999; Markow and O’Grady 2005; Laturney and Billeter 2014; Anholt, et al. 2020). However, while there is strong evidence that links sex differences in expression with sex dimorphism, the underlying mechanisms of species differences in sex dimorphism are poorly understood. A study by Graze, et al. 2012 (Graze, et al. 2012) found that genes with *D. simulans*-biased alleles in interspecific hybrids were enriched for genes associated with the GO term “H3K4 methyltransferase activity” and *D. melanogaster-biased* alleles to be enriched for genes associated with the GO term “H3K9 methyltransferase activity”. H3K9 methylation is correlated with closed chromatin and silenced expression (reviewed in Boros 2012; Kimura 2013), similar to H3K27 methylation. This finding led us to hypothesize that there may be divergence in chromatin patterns between the species.

Sex-biased gene expression in brain, eye, and antennal genes has been shown to be associated with sexually dimorphic behavior and sensory perception (Landry, et al. 2007; Kopp, et al. 2008; Shiao, et al. 2015; reviewed in Anholt, et al. 2020). Sex-biased gene expression, or a gene with greater expression in one sex over the other, has been previously shown to be rapidly evolving (e.g., Ellegren and Parsch 2007; Zhang, et al. 2007; Harrison, et al. 2015). Studying the evolution of gene expression regulation within a relatively short period of evolutionary time, such as between two closely related species, allows for identification of the sets of genes, mechanisms and processes contributing to speciation and the evolution of species differences. *D. melanogaster* and *D. simulans* have diverged relatively recently in evolutionary history (~5 million years ago) (Tamura, et al. 2004) and *D. melanogaster* and *D. simulans* have diverged in many sexually dimorphic phenotypes, including courtship behavior (Cobb, et al. 1989). Additionally, hybrid studies of *D. melanogaster* and *D. simulans* have suggested divergence of sex-biased expression regulatory mechanisms between the species (Ranz, et al. 2004). In combination with the wealth of resources available for Drosophila as a model organism, the comparison of *D. melanogaster* and *D. simulans* provides an exceptionally tractable model in which to explore the relationship between chromatin marks and sex difference in expression, in an evolutionary context. To this end, we assess the relationship between sex-biased expression and chromatin accessibility within each species, as well as how this relationship evolves in two closely related species.

## Results

We assayed males and females in the sister species, *D. melanogaster* and *D. simulans*, for gene expression (n=48; 2 sexes x 2 genotypes x 2 species x 6 replicates), and chromatin (n=24; 2 sexes x 1 genotypes x 2 species x 6 replicates). For each sample ChIP for the open chromatin mark, H3K4me3, and closed chromatin marks, H3K27me2me3 and input were collected. We compared the two sexes within each species, trends of sex-bias between species, and one-to-one orthologous loci between species for gene expression and chromatin and evaluated the relationship between sex-bias in gene expression and chromatin status. Within *D. melanogaster*, 2,556 genes on the X chromosome and 14,114 genes on the autosomes were examined, and in *D. simulans*, 2,305 genes on the X and 12,504 genes on the autosomes. There were 1,840 X and 10,097 autosomal one-to-one orthologs used to compare species gene-to-gene. We performed extensive quality control of the data (See Supplementary Materials Sections 4-7). For example, to evaluate whether genome quality affected the results all analyses were also performed with both species mapped to *D. melanogaster* (FlyBase r6.17) and both species mapped to *D. simulans* (FlyBase r2.02). While there were a few genes with consistent map bias, there was no evidence that genome quality impacted mapping (mapping rates were similar between species) and no trends reported were affected by the choice to map each species to its own genome rather than mapping both to one of the two genomes (Supplementary Materials Section 5.3).

Exonic regions were separated into non-overlapping exonic features where alternative donor/acceptor sites were quantified separately from shared exonic regions, in order to capture the potential sex-specific exonic features in each gene (Newman 2018). Non-overlapping exonic features were quantified as *C_is_* = (∑(*d_ijs_*)/*N_i_*) × (*Q*/*U_s_*), where *d* is the depth of reads at nucleotide *j* of feature *i*, *N* is the length of the feature, *U_s_* is the upper quartile of (∑(*d_ijs_*)/*N_i_*) values in sample s, and *Q* is the median of all *U_s_* values within the given species (Bullard, et al. 2010; Dillies, et al. 2013).

If all detected features were detected in only one sex, the gene was labeled as sex-limited. There were 770 genes (~6% of expressed genes) determined to be sex-limited in *D. melanogaster* (569 in males, 201 in females) and 547 genes (~4% of expressed genes) in *D. simulans* (352 in males, 195 in females) (Supplementary File 1, Supplementary File 2, *flag_sex_limited*==1). Differential expression analyses were performed separately for each exonic feature detected in both sexes of each species. Genes were considered sex-biased in expression if at least one exonic feature was statistically significantly differentially expressed between sexes. Genes with both significantly male- and female-biased exonic features were designated “Male-biased and Female-biased” and are expected in genes that are sex-specifically alternatively spliced, such as the sex determination gene *dsx* (Supplementary Figure 1).

ChIP samples were compared to input controls for genomic features (Transcription start sites, 5’, 3’ UTR’s, exonic features and introns). Genomic features were considered detected above the input control in H3K4me3 (DAI) if *C*_*K*4,*is*_ > *C_Input,is_*, in more than 50% of the replicates for that species-sex combination, and as *C*_*K*27,*is*_ > *C_Input,is_*, for H3K27me3me4. ChIP data were found to be high quality and conform with general expectations for detection of the marks (Supplementary Materials Sections 7.1-7.2). A gene was considered as having a mark if at least one exonic feature in the gene was DAI. A gene was considered sex-limited (male/female) when marks were detected in only one sex.

### Genes that diverge in sex-bias between the species are more likely to be male-biased than female-biased

Gene expression in head tissues was measured in independent replicates of males and females for each species (n = 48, 2 species x 2 sexes x 2 genotypes x 6 replicates). In addition to the excess of male-limited expression compared to female limited expression, there is an excess of male-biased expression compared to female-biased expression observed in both *D. melanogaster* (2723 male-biased vs. 2185 female-biased, Binomial *p* < 0.0001) and *D. simulans* (2160 male-biased vs. 1873 female-biased, Binomial *p* < 0.0001) is statistically significantly different in orthologous genes where sex-bias is gained/lost between species (Table 1, rows 13 vs. 14 and 16 vs. 17, Supplementary Figure 2C). Although the number of male biased orthologs was greater than the number of female biased orthologs (Table 1, rows 1 vs. 2), there was no statistically significant excess of male-bias based on our threshold of *p* < 0.001. For orthologs unbiased in *D. melanogaster* and sex-biased in *D. simulans*, more genes were male-biased than female-biased (Binomial *p* = 0.0001, Table 1, Supplementary Figure 2C). Similarly, for orthologs unbiased in *D. simulans* and sex-biased in *D. melanogaster*, we found an excess of male-biased expression compared to female-biased expression (Binomial *p* < 0.0001, Table 1, Supplementary Figure 2C).

**Table 1.**
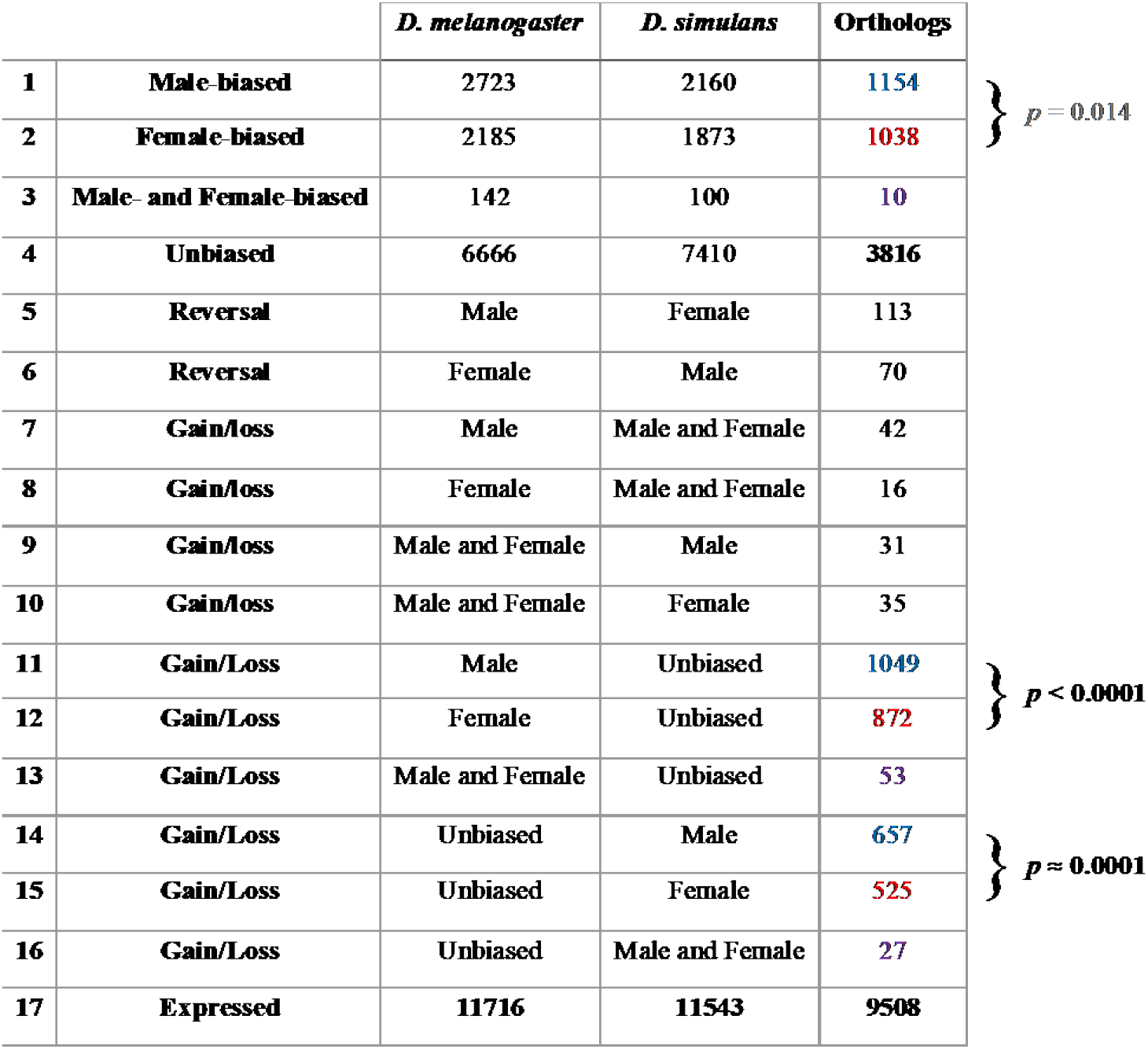
Sex bias in expression. The observed pattern of sex-bias is listed in column 2, all definitions are mutually exclusive. In rows 1-4 the number of genes following the pattern in column 1 for each species are given in columns 3 and 4 and the number of orthologous that have the same pattern in both species are in the right-most column. Larger numbers in *D. simulans* and *D. melanogaster* columns reflect observations for genes for which no one-to-one ortholog was identified or the one-to-one ortholog was not expressed in both sexes of both species. There are a total of 5050 sex-biased genes in *D. melanogaster*, 4133 in *D. simulans*, and 2202 orthologs with the same sex-bias observed in both species. Rows 5-17 are the observed patterns of sex bias where the two species diverge. Binomial test probabilities are indicated to the right of the table for the comparison of male-biased vs. female-biased for consistent and species-specific sex-biased gene classifications. P-values are in black if below the significant threshold of *p* = 0.001 and gray if above the threshold. Reversal of sex-bias is rare, only two percent (183 / 9508) of orthologs. Genes on chromosome 4 and on scaffolds, as well as those that change location are omitted. Values of the X and autosomes separately for each category are listed in Supplementary Table 1.

### Male-biased orthologs are less constrained than female-biased orthologs

The estimated sex-ratio in orthologs is strikingly concordant for females (β_1f_ = 0.6428; Pearson’s *r* = 0.69; T-test *H*_0_: β_1_ = 0, *p* < 0.0001) and males (β_1_ = 0.4663; Pearson’s *r* = 0.49; T-test *H*_0_: β_1m_ = 0, *p* < 0.0001). Male-biased orthologs, are less concordant than female-biased orthologs (*H*_0_: β_1m_ = β_1f_, *p* < 0.0001) (Figure 1A). In addition, since the regression coefficients of both male-biased orthologs and female-biased orthologs are significantly less than 1 (T-test *H*_0_: β_1_ ≥ 1, *p* < 0.0001 for both male-biased and female-biased orthologs), there is evidence for a larger sex-bias in *D. melanogaster* compared to *D. simulans*. Orthologs with gains/losses in sexbias show large variation in the magnitude of the ratio of sex-bias compared to those with male or female bias (Figure 1B).

**Figure 1.**
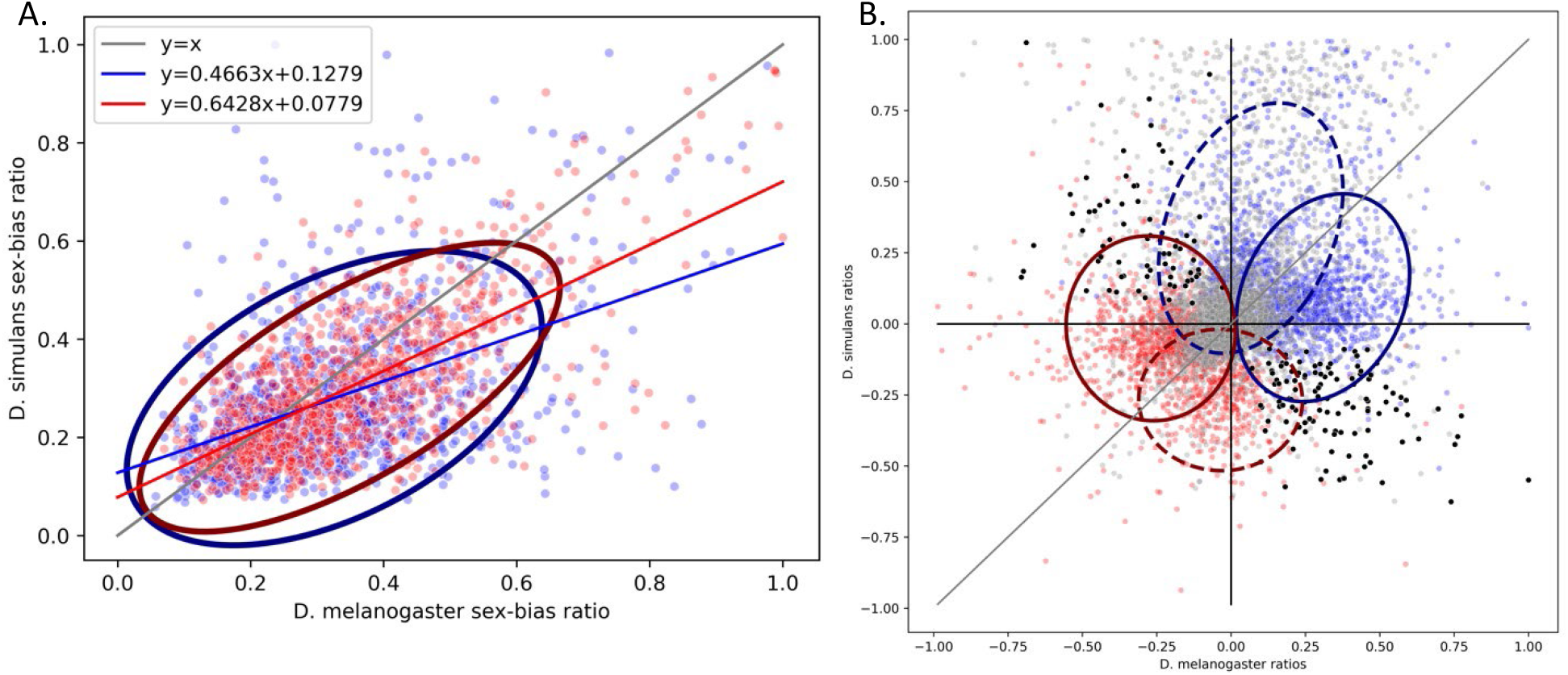
Sex bias ratios across orthologs. (Panel A) For the orthologous genes where sex bias is in the same direction between the two species, we examined the relationship between the observed ratio of sex bias in *D. melanogaster* (X-axis) and *D. simulans* (Y-axis). The Y=X line is in gray. To compare female-biased and male-biased genes on the same plot we calculated the sexbias ratio here as 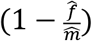 for male-biased orthologs (blue dots), and 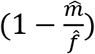 for female-biased orthologs (red dots); where 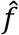 is the average UQ normalized expression across female samples and *m* is the average UQ normalized expression across male samples, A value close to 1 indicates extreme sex-bias, while a value close to 0 indicates low sex-bias. A linear regression of the *D. melanogaster* estimate on the *D. simulans* estimate was calculated for female-biased (red) or male-biased (blue) orthologs separately. The ellipses represent the 95^th^ percentile of the observed data. (Panel B) Gains and losses in sex-bias. *D. melanogaster* sex-bias ratios (X-axis) compared to *D. simulans* sex-bias ratios (Y-axis). In order to visually separate the male and female bias, we calculated the sex-bias ratio as 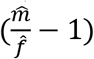 if 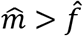, and as 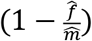 if 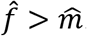. Orthologs significant for male-bias in one species are colored blue, and those significant for female-bias in one species are colored red. The solid ellipses represent the 95^th^ percentile of the observed statistically significant species-specific sex bias in *D. melanogaster*. The dashed ellipses represent 95^th^ percentile of the observed statistically significant species-specific sex bias in *D. simulans*. Orthologs with no significant sex-bias in either species are plotted in gray. Orthologs with reversal of sex bias are potted in black (n=183, 2% of all orthologs).

### Male-bias in orthologs is associated with signatures of positive selection

The comparative genomics database, flyDIVas (Clark, et al. 2007; Stanley and Kulathinal 2016) provides gene-level estimates of divergence with nonsynonymous (dN) to synonymous substitution (dS) rates (dN/dS) and tests of positive selection using PAML (Yang 1997) for the melanogaster subgroup (*D. melanogaster, D. simulans, D. sechellia, D. yakuba, D. erecta*), melanogaster group (melanogaster subgroup and *D. ananassae*), and the 12 Drosophila species (melanogaster group and *D. pseudoobscura, D. persimilis, D. willistoni, D. mojavensis, D. virilis*, and *D. grimshawĩ*). These different group allow for evaluation of selection across these three levels of phylogenic depth; however, the number of orthologous loci does decline as the distance from melanogaster increases and the tests at the 12 genome level are to be thought of as suggestive (Stanley and Kulathinal 2016). The null hypotheses tested are codon-based tests of positive Darwinian selection based on d_N_d_s_ (ω) ratios estimated by PAML model M0 (Yang 1997). Three nested pairs of site-specific models are available on flyDIVas: 1) model M1a (neutral) vs. M2a (positive selection), 2) model M7 (beta-distributed) vs. M8 (beta+ω>1) (Yang 1997), and 3) model M8 (beta+ω>1) vs. model M8a (beta+ω=1) (Swanson, et al. 2003; Wong, et al. 2004). This is a set of 9 tests of association (Supplementary Table 2). We used *p* < 0.001 as the significance threshold and find that only the *D. melanogaster-D. simulans* male-biased orthologs are significantly enriched for positive selection. This is a consistent inference for 8 of the 9 tests with the exception being test M8 vs. M8a in the 12 species comparison (χ^2^: *p* = 0.10) (Supplementary Table 2). There were 6 genes where reversals in sex bias occurred where individual genes showed signatures of positive selection (*Exn, yin, SNF4Aγ, Esyt2, DIP*-η, and *milt*).

### Sex-bias is enriched on the X chromosome

Sex-biased expression in orthologs is enriched on the X chromosome compared to the autosomes (Fisher’s exact test: overall sex-biased *p* < 0.0001, male-biased *p* < 0.0001, female-biased *p* < 0.0001). No significant enrichment on the X is observed in the genes with gains/losses of sex bias between the species according to our threshold of *p* < 0.001 (Supplementary Figure 3), including orthologs sex-biased in *D. melanogaster* and unbiased in *D. simulans* (Fisher’s exact test: overall sex-biased *p* = 0.002, male-biased *p* = 0.05, female-biased *p* = 0.03) and orthologs sex-biased in *D. simulans* and unbiased in *D. melanogaster* (Fisher’s exact test: overall sex-biased *p* = 0.43, male-biased *p* = 0.42, female-biased *p* = 0.54). Although, there were more orthologous genes with sex-biased expression in *D. melanogaster* compared to *D. simulans* (Table 1, rows 13-15 vs. 16-18; McNemar: *p* < 0.0001). These results are robust to map bias (See Supplementary Materials Section 5.3).

### Genic chromatin marks are conserved

The marginal frequencies for both open and closed chromatin differ from 0.5. In order to evaluate the agreement in marks, we use a chance corrected measure of agreement (kappa, κ) to account for these differences in the marginal frequencies (Fleiss 1981) (Supplementary Figure 4). A κ =1 indicates perfect agreement, and negative values indicate less agreement than expected by chance and a value of 0 indicates no agreement. Agreement between sexes and species is high for both chromatin marks (Supplementary Table 3). Within each species, agreement between males and females for the presence of H3K4me3 was high (*D. melanogaster* κ = 0.73, *D. simulans* κ = 0.68), as well as for H3K27me2me3 (*D. melanogaster* κ = 0.58, *D. simulans* κ = 0.54). Additionally, agreement between the species is high for H3K4me3 in males (κ = 0.67) and females (κ = 0.73) and for H3K27me2me3 in males (κ = 0.52) and females (κ = 0.54). There is a set of 7,714 orthologs (~65% of all one-to-one orthologs) with H3K4me3 marks present in both sexes and both species and 1,817 (~16%) orthologs with H3K27me2me3 present in both sexes and both species. These common marks make up ~76% of all genes with H3K4me3 and ~30% of genes with H3K27me2me3. However, as expected, there are very few genes with both marks. The agreement between genes with H3K4me3 and H3K27me2me3 is negative for both sexes and both species indicating that these marks coincide less frequently than expected by chance (Supplementary Table 3).

### Sex-limited chromatin accessibility diverges

We find there is more open chromatin in *D. simulans* compared to *D. melanogaster* (McNemar: *p* < < 0.0001) and more closed chromatin in *D. melanogaster* compared to *D. simulans* (McNemar: *p* < < 0.0001). While agreement between species is high in marks overall, there is low agreement when marks are sex-limited (H3K4me3 κ: 0.05-0.30, H3K27me2me3 κ: 0.05-0.15, Supplementary Table 3). However, there are more genes with conserved male-limited H3K4me3 than female-limited H3K4me3 on the X (Binomial *p* < 0.0001, Figure 2B) and autosomes (Binomial *p* < 0.0001, Figure 2C), while there are nearly equal numbers of genes with conserved male-limited and female-limited H3K27me2me3 on both the X (Binomial *p* = 0.002, Figure 2B) and autosomes (Binomial *p* = 0.83, Figure 2C). Interestingly, on the autosomes, sex-limited H3K4me3 shows more genes with male-limited marks in *D. simulans* compared to *D. melanogaster* (McNemar: *p* < < 0.0001, Figure 2A) and female-limited marks are more prevalent in *D. melanogaster* compared to *D. simulans* (McNemar: *p* < 0.0001, Figure 2A).

**Figure 2.**
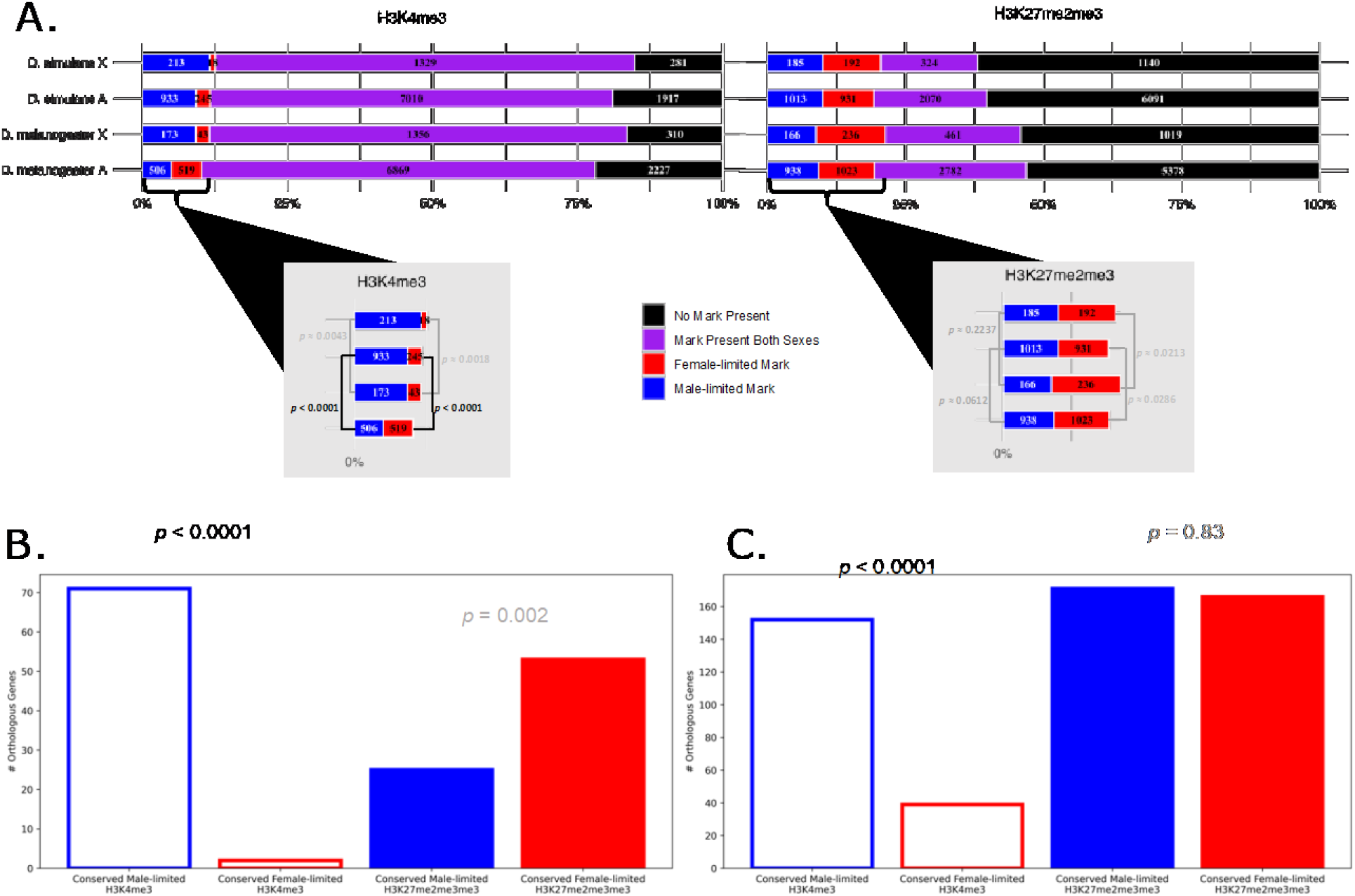
Chromatin marks in *D. melanogaster* and *D. simulans*. The number of orthologs (n=12,083) with male-limited, female-limited, or marks in both sexes indicated in blue, red, and purple respectively. Most marks are detected in both sexes. Panel A) In *D. melanogaster*, 1,882 and 10,121 genes are on the X and autosomes respectively, and 1,841 and 10,105 for *D. simulans* X and autosomes. There are 1,840 genes on the X of both species, 10,097 genes on the autosomes of both species, 7 genes on the X of *D. melanogaster* and autosomes of *D. simulans*, and 1 gene on the X of *D. simulans* and autosomes of *D. melanogaster*. The differences between the presence of marks in males compared to females was evaluated using McNemar test (McNemar 1947) with p-values for each test indicated in black for significant (< 0.001) and gray otherwise. Genes on the X (Panel B) and autosomes (Panel C) with conserved male-limited and female-limited chromatin marks (where both species are sex-limited in the same direction for a given chromatin mark in a gene) are indicated in blue or red for male-limited and female-limited respectively, and open box for the H3K4me3 and filled box for H3K27me2me3. P-values for binomial tests between the number of genes with male-limited and female-limited are indicated in black for significant (< 0.001) and gray otherwise.

### Chromosomal bias in chromatin

There is a higher proportion of genes with open chromatin marks detected in both species on the X chromosome compared to the autosomes (Supplementary Figure 5, Supplementary Figure 6D). We observe more male-limited than female-limited H3K4me3 on the X chromosome of *D. simulans* (McNemar: *p* < 0.0001) and *D. melanogaster* (McNemar: *p* < 0.0001) and conserved male-limited H3K4me3 marks are enriched on the X compared to the autosomes (χ^2^: *p* < 0.0001) while conserved female-limited H3K4me3 marks have no chromosomal bias (Fisher exact: *p* = 0.08). Concomitantly, genes with conserved presence of female-limited H3K27me2me3 marks are enriched on the X compared to the autosomes (χ^2^: *p* = 0.0004) and no chromosomal bias is observed for male-limited H3K27me2me3 marks (χ^2^: *p* = 0.35).

### Sex-biased expression is associated with open chromatin

We propose a model, “Open in Same sex and/or Closed in Opposite” (OS-CO), as an expectation of chromatin accessibility states for genes with sex-biased expression (Figure 3). We expect chromatin in male-biased genes to have i) open chromatin marks in males, and/or ii) closed chromatin marks in females. We test this expectation by comparing the chromatin state in male-biased genes to genes without male bias using Fisher exact test (Fisher 1934). Under the null hypothesis that chromatin is independent of sex-bias there should be no difference in the proportion of genes with open chromatin in males in these two groups. Similarly, we compare the presence of open chromatin marks in females between female biased genes and non-female biased genes. In both species, open chromatin marks in females are more likely to occur in female-biased genes relative to non-female-biased genes (*D. melanogaster* χ^2^: *p* < 0.0001; *D. simulans* χ^2^: *p* < 0.0001; Figure 4A) and open chromatin marks in males are enriched in genes with male expression bias compared to genes without male bias in expression (*D. melanogaster* χ^2^: *p* < 0.0001; *D. simulans* χ^2^: *p* < 0.0001; Figure 4B).

When comparing the X and autosomes separately, female-biased genes showed the same pattern of association with chromatin marks as in the combined set of genes across the genome (Figure 4C). Genes with male-biased expression were enriched for female closed chromatin on the autosomes in both *D. melanogaster* and *D. simulans*, but on the X chromosome the chromatin pattern was divergent between the two species. In *D. melanogaster* there was an enrichment for male-biased expression with female closed chromatin on the X, whereas in *D. simulans* there was not (Figure 4D). Outside of this divergence on the X, the expression and chromatin association patterns are remarkably similar between the species on both the X and autosomes, with the most striking differences observed between the sexes.

**Figure 3.**
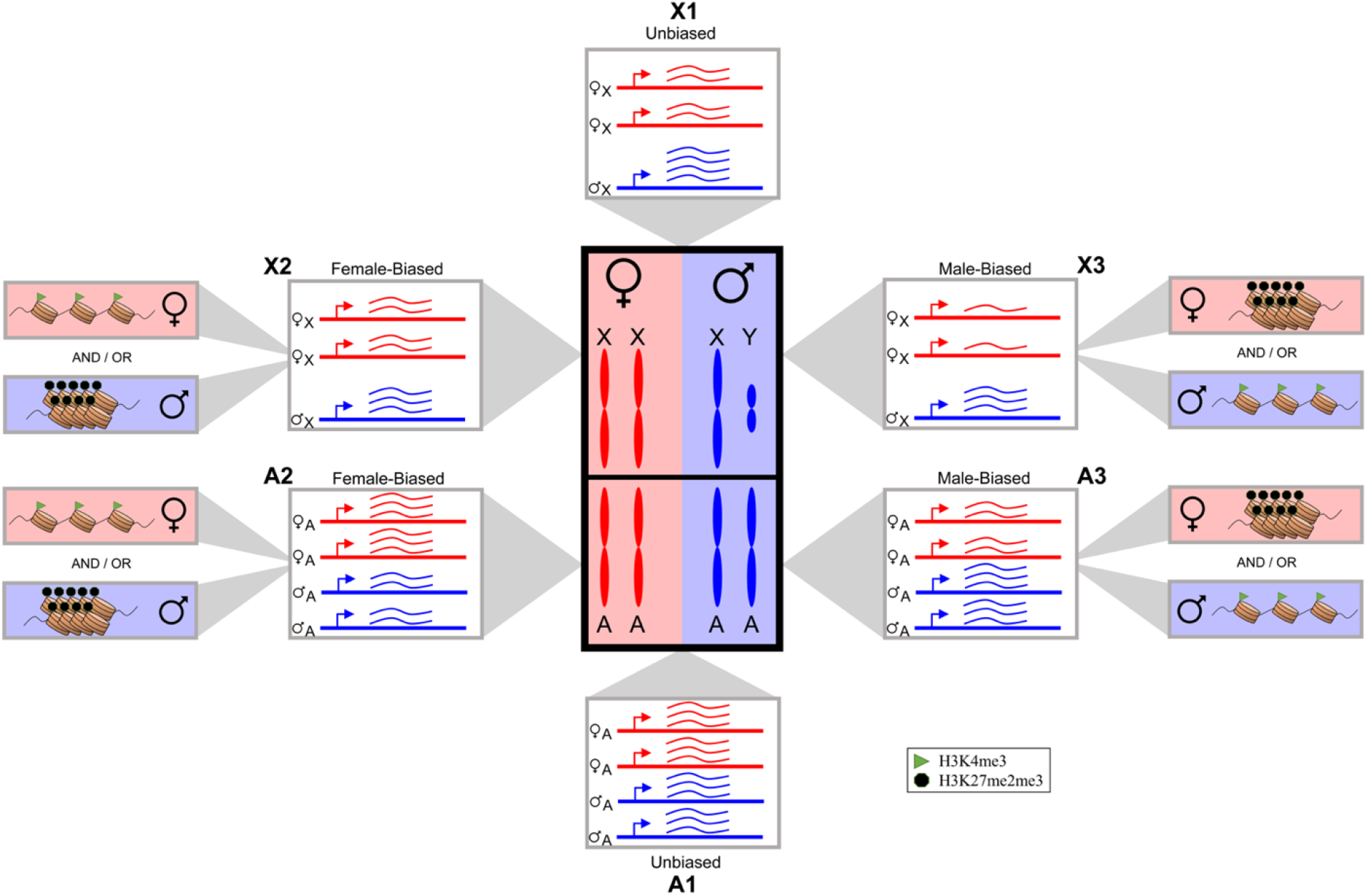
“Open in Same and/or Closed in Opposite” (OS-CO): a model for chromatin accessibility patterns for sex-biased expression. Representation of gene expression categories between males and females on the X chromosome (X) and autosomes (A). Unbiased (X1, A1) genes are defined as those without statistical evidence of differential expression. Female-biased (X2, A2) genes are those with at least one exon with statistical evidence towards female expression. Male-biased (X3, A3) genes are similarly defined towards male expression. Female-biased (X2, A2) expression patterns are expected to have open chromatin marks (H3K4me3) in females and/or closed chromatin marks (H3K27me2me3) in males. The mirror pattern is expected for male-biased (X3, A3) expression patterns are expected to have open chromatin marks (H3K4me3) in males and/or closed chromatin marks (H3K27me2me3). Not all sex-biased genes are expected to have these patterns as there are other chromatin marks and regulatory factors that may influence expression.

**Figure 4.**
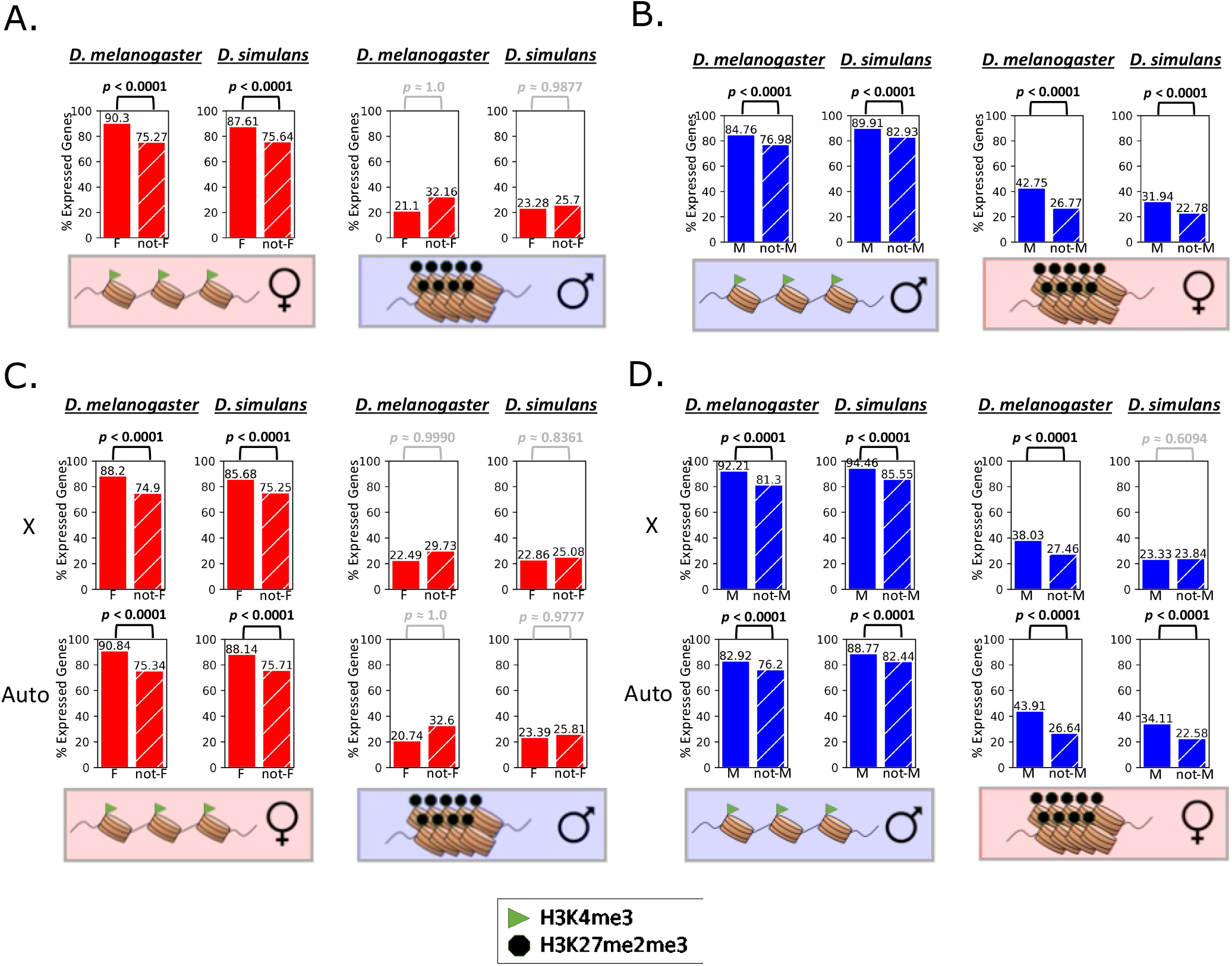
Sex-biased expression is associated with chromatin marks. The Y-axis of each graph represents the percent of expressed female-biased (solid red), non-female-biased (hatched red), male-biased (solid blue), or non-male-biased (hatched blue) genes within each species with the indicated chromatin (cartoon representations below each set of bars). Consistent with the model presented in Figure 4, (Panel A) Female-biased genes (solid red) are enriched for H3K4me3 (open) chromatin when compared to non-female-biased genes (hatched red) in both species. (Panel B) Male-biased genes (solid blue) are enriched for male open chromatin and female H3K27me2me3 (closed) chromatin when compared to non-male-biased genes (hatched blue) in both species. The model in Figure 4 was also evaluated for X and autosomes separately. (Panel C) Female-biased genes (solid red) are enriched for open chromatin when compared to non-female-biased genes (hatched red) on both the X and autosomes of both species. (Panel D) Male-biased genes (solid blue) are enriched for male open chromatin and female closed chromatin when compared to non-male-biased genes (hatched blue) on both the X and autosomes of *D melanogaster*. *D simulans* shows the same pattern on the autosomes. On the X chromosome, male-bias genes are enriched for open chromatin in males but not for closed chromatin in females, showing a divergence in the regulatory pattern between the two species. There were 11,716 (n_X_=1,919, n_A_=9,797) genes expressed in *D. melanogaster* and 9,902 genes expressed in *D. simulans* (n_X_=1,893, n_A_=9,650) evaluated for sex-biased expression and chromatin presence. Each set of female-biased (male-biased) and non-female-biased (non-male-biased) genes were tested for enrichment of the indicated chromatin mark using Fisher exact test (Fisher 1934) with the alternative expectation that the indicated chromatin marks would be more likely in genes with female-biased (male-biased) expression. Significant p-values (*p* < 0.001) are black and p-values above the significance threshold are gray.

There are 34 genes on the X chromosome with male-biased expression in both species but female closed chromatin only in *D. melanogaster* and not *D. simulans*. These genes are contributing to the different patterns of chromatin mark usage observed on the X chromosomes of the two species in Figure 4D. These 34 genes include the well described *D. melanogaster-D. simulans* hybrid incompatibility gene, *Hmr* (Hutter and Ashburner 1987; Barbash, et al. 2003) that also has been associated with heterochromatin factors (Satyaki, et al. 2014). Genes associated with habituation [*wcy*, (Lugtenberg, et al. 2016)] and behavior [*Adar*, (Palladino, et al. 2000); *norpA*, (Pick and Strauss 2005)] were also observed in this set of 34 genes. Further study of the *D. melanogaster-* specific sex-biased genes with divergent chromatin regulation may reveal insights into sex-dependent gene expression evolution and the role chromatin accessibility may play in the evolution of these genes.

### Sex-biased orthologs have conserved presence of open chromatin

In both species, the vast majority sex-biased orthologs have open chromatin in the sex with greater expression (Figure 4, Supplementary Figure 7) consisted with our model (Figure 3). Male-biased orthologs are significantly enriched for conserved open marks in males (~89% of male-biased orthologs vs ~76% of unbiased orthologs; χ^2^: *p* < 0.0001). Similarly, female-biased orthologs are significantly enriched for conserved open marks in females (~90% of female-biased orthologs vs ~73% of unbiased orthologs; χ^2^: *p* < 0.0001). In addition, the agreement for H3K4me3 marks within species in male-biased orthologs (*D. melanogaster* 0.63; *D. simulans* 0.60) and female biased orthologs (*D. melanogaster* 0.65; *D. simulans* 0.60), is lower than for unbiased orthologs (*D. melanogaster* 0.71; *D. simulans* 0.66). When male-biased orthologs have conserved H3K4me3 marks in males and no H3K27me2me3 mark in females, the sex-bias ratio is more similar (Figure 5A; β_1_ = 0.5023) than when both species have a female H3K27me2me3 mark (Figure 5C; β_1_ = 0.1795; β_1A_ vs. β_1C_: *p* < 0.0001). When the female mark is not conserved between the species (present in either species the sex-bias ratio is more conserved than when both species have the female mark (Figure 5B; β_1_ = 0.4099; β_1A_ vs. β_1B_: *p* ≈ 0.1590) (for all combinations of chromatin marks for male-biased orthologs see Supplementary Figure 8). In contrast, the sex-ratio for female-biased orthologs does not change with the male H3K27me2me3 chromatin mark (Figure 4, Figure 5D-F, Supplementary Figure 9).

**Figure 5.**
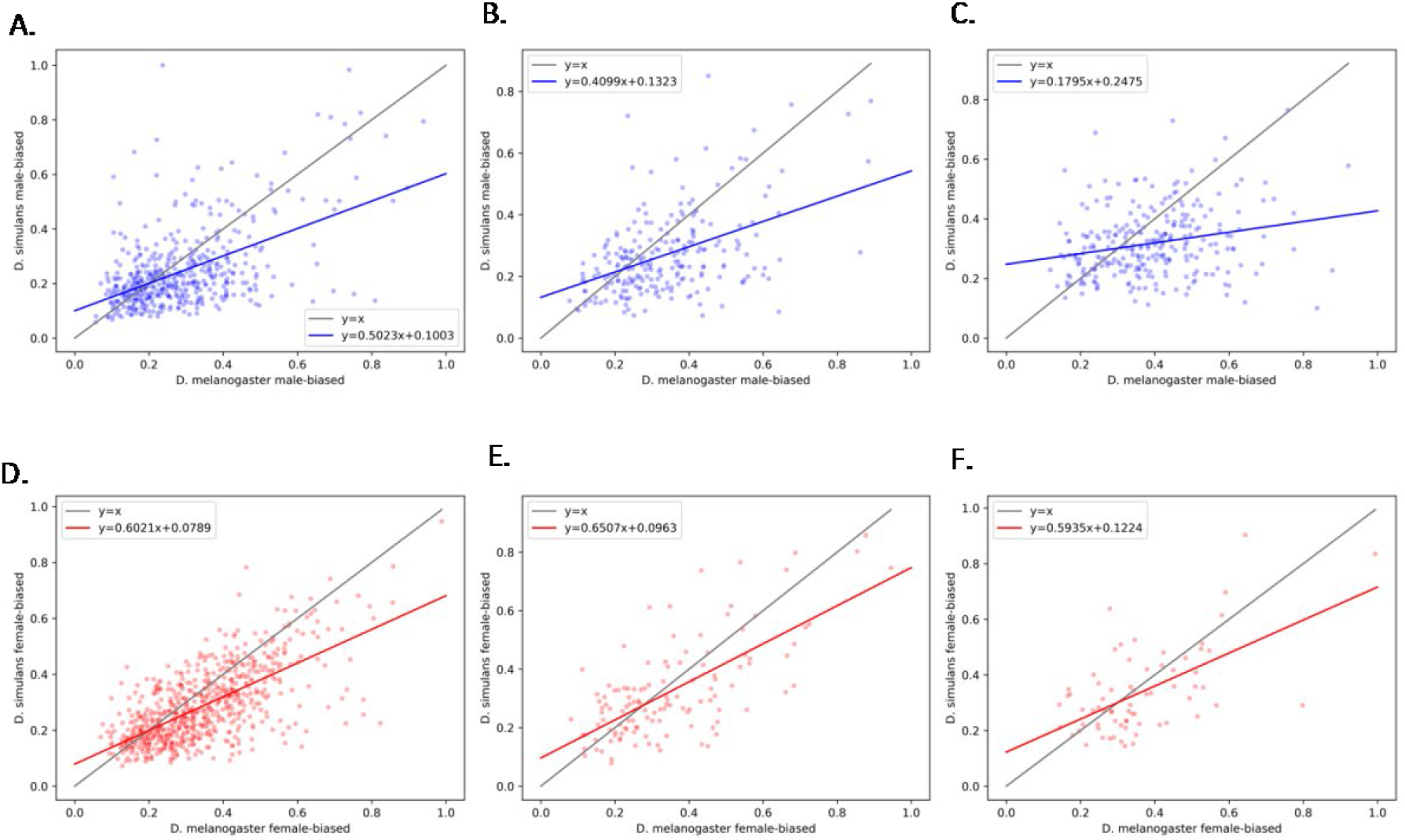
Male and female biased orthologs. Estimated ratio of sex-bias in *D. melanogaster* (X-axis) and *D. simulans* (Y-axis), with the y=x line in gray. To compare female-biased and male-biased genes on the same scale, (0,1) we plotted 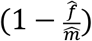 for male-biased orthologs (blue dots), where 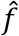 is average UQ normalized expression across female samples and 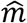 is average UQ normalized expression across male, and 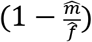 for female-biased orthologs (red dots). Genes are separated by chromatin presence within the species. For male-biased orthologs: (Panel A) conserved presence of male H3K4me3 and absence of female H3K27me2me3, (Panel B) conserved presence of male H3K4me3 and female H3K27me2me3 in one species only, and (Panel C) conserved presence of male H3K4me3 and female H3K27me2me3. For female-biased orthologs: (Panel D) conserved presence of female H3K4me3 and absence of male H3K27me2me3, (Panel E) conserved presence of female H3K4me3 and male H3K27me2me3 in one species only, and (Panel F) conserved presence of female H3K4me3 and male H3K27me2me3. A red or blue line indicates the linear regression calculated for the conserved female-biased or conserved male-biased genes respectively using the least-squares method. Regression coefficients for each panel are as follows: Panel A β_1_ = 0.5023, Panel B β_1_ = 0.4099, Panel C β_1_ = 0.1795, Panel D β_1_ = 0.6021, Panel E β_1_ = 0.6507, and Panel F β_1_ = 0.5935.

## Discussion

We propose a new model of how chromatin state is associated with sex-bias in expression. This model hypothesizes that in genes with male-biased expression we expect to see an excess of open chromatin in males compared to genes without male-bias; and in genes with female-biased expression we expect to see an excess of open chromatin in females compared to genes without female-bias. That is, we expect to observe open chromatin in the sex where we see expression bias (OS). We also hypothesize that sex-bias in expression toward one sex may be associated with closed chromatin in the opposite sex (CO). While, the open chromatin mark H3K4me3 is correlated with active expression (Santos-Rosa, et al. 2002; Schneider, et al. 2004) and the closed chromatin marks H3K27me2 and H3K27me3, together referred to as H3K27me2me3, are correlated with silenced expression (Wang, et al. 2008; Juan, et al. 2016); these marks do not act independently to affect chromatin accessibility. In Drosophila embryos, expression variation has been found to be more predictive of the open chromatin mark H3K4me3 rather than the reverse (Floc’hlay, et al. 2021), supporting the hypothesis that H3K4me3 does not induce transcription but is instead deposited as a result of active transcription (reviewed in Howe, et al. 2017). Other histone modifications, DNA methylation, and chromatin factors are involved in the establishment and plasticity of chromatin accessibility (reviewed in Boros 2012). It is likely that our observations using these marks does not completely reflect final active or repressed states of expression resulting from the chromatin state as a whole, as we assayed only 2 of the many possible marks. Our study does not demonstrate a causal relationship between chromatin accessibility and sex-biased expression, nor do we claim to provide a comprehensive survey of chromatin accessibility. Rather, our findings likely reflect the role of different regulators that impact chromatin states. Even with broad limitations with respect to the suite of marks assessed, the OS component of the model holds broadly. *Genes with sex-biased expression are more likely to have H3K4me2 marks in the sex with greater expression in D. melanogaster and D. simulans, for both sexes, on both X and autosomes compared to unbiased genes*.

The direction of sex-bias in expression agreed between the two species much more frequently than expected by chance (male-bias: κ = 0.41, *p* < 0.0001; female bias: κ = 0.45, *p* < 0.0001). This agreement in presence/absence of sex-bias between *D. melanogaster* and *D. simulans* may be due to the short evolutionary time and the maintenance of the ancestral state where the sex-bias in the common ancestor is random. Consistent with drift, the proportion of orthologs with male-bias is not different from those with female-bias at our threshold (*p* < 0.001), although the number of male biased orthologs is greater than the number of female biased orthologs. Under the null hypothesis that the direction of bias is random, we would also expect to see approximately an even number of gains/losses in transitions between the two species from unbiased to male- or female-biased. In a binomial test, the null hypothesis of equal probability for male/female gain/loss (p=0.5) is rejected for both transitions from unbiased genes in *D. melanogaster* to sex biased genes in *D. simulans* (~55% male-biased, Binomial *p* < 0.0001) and unbiased genes in *D. simulans* to sex-biased genes in *D. melanogaster* (~56% male-biased, Binomial *p* ≈ 0.0001). There is also more male-bias than female-bias in sex limited expression (*p* < 0.0001 for both species).

Sex-bias is conserved in magnitude, as well as direction. Intriguingly, sex-bias ratios for expression are more similar between the species in females than males, suggesting there may be either less constraint in males, or potentially a difference in selection between the sexes. While there is no evidence in female-biased orthologs for the CO portion of our model, in male-biased orthologs, the magnitude of sex-bias is affected by the presence of the female closed chromatin marks providing some support for this hypothesis.

The excess of male-bias in sex-limited gene expression in both species, coupled with a significant excess of male-bias in orthologs in the gain/loss of sex-bias, and less conservation in the magnitude of the sex-bias ratio suggests that there is a possibility that the male-biased genes are evolving faster. Male-biased genes have been shown to be evolving faster than other genes in comparisons between *D. melanogaster* and *D. simulans* (Meiklejohn, et al. 2003) with overall higher rates of evolution in male-biased genes observed in gonadal tissue (Perry, et al. 2014; Whittle and Extavour 2019) as well as whole body or somatic tissue (Ranz, et al. 2003; Zhang, et al. 2004; Connallon and Knowles 2005; Ellegren and Parsch 2007).

In previous studies of D. melanogaster, head and brain tissues have been reported to have more male-biased than female-biased expression (Chang, et al. 2011; Catalan, et al. 2012; Newell, et al. 2016; Palmateer, et al. 2021) with enrichment for male-biased genes on the X chromosome compared to the autosomes (Goldman and Arbeitman 2007; Chang, et al. 2011; Catalan, et al. 2012; Meisel, et al. 2012a; Huylmans and Parsch 2015). Whole body tissue has been observed to have more female-biased expression than male-biased expression (Ranz, et al. 2003; McIntyre, et al. 2006; Wayne, et al. 2007; Graze, et al. 2014; Allen, et al. 2017) and enrichment for female-biased genes on the X compared to the autosomes (Ranz, et al. 2003; McIntyre, et al. 2006; Wayne, et al. 2007; Meisel, et al. 2012a; Graze, et al. 2014). We do find some evidence of positive selection in male-biased orthologs. However, we cannot exclude the possibility that we observe an excess in male-bias due to a relaxation of constraints in this specific tissue. The smaller slope in the comparison between the species of the magnitude of sex-bias ratios in male-biased orthologs compared to female-biased orthologs supports the relaxation of constraint hypothesis.

As terminal transcription factors of the sex determination pathway, *dsx* and *fru* have male- and female-specific isoforms (Supplementary Figure 1). *Dsx* contributes to the regulation of sexual dimorphism in the brain of both sexes (Rideout, et al. 2007; Kimura, et al. 2008; Rideout, et al. 2010; Arbeitman, et al. 2016), and is conserved among Drosophila species (Shukla and Nagaraju 2010). Although female-biased orthologs were not enriched for genes regulated by *dsx* (Arbeitman, et al. 2016)(χ^2^: *p* = 0.664), male-biased orthologs were enriched for genes regulated by dsx (χ^2^: *p* < 0.0001). *Fru*, is highly conserved in sex-specific splicing across insects (Salvemini, et al. 2010). Fru^M^ is associated with chromatin remodeling factors (Lorbeck, et al. 2010; Ito, et al. 2012). Additionally, the *fru* gene itself may be regulated by pheromone-induced chromatin remodeling at the *fru* promotor in specific neurons (Zhao, et al. 2020) and *fru* expression decreases with mutation in histone demethylase *Kdm4A* (Lorbeck, et al. 2010). Expression of Fru^M^ has been shown to affect the establishment of closed chromatin marks in male neurons (Brovkina, et al. 2021) resulting in the repression of genes that lead to sex-specific phenotypes (Ito, et al. 2016; Sato, et al. 2020; reviewed in Goodwin and Hobert 2021). Palmateer et al. show overall differences in chromatin patterns within genes enriched in *fru-P1* TRAP experiments (Palmateer, et al. 2021). Consistent with the potential role of Fru^M^ as an activator of expression in males (Vernes 2014) is the excess of open chromatin in males compared to female-limited open chromatin for both species in this study on the X and for *D. simulans* on the autosomes. The male-specific Fru^M^ protein is a primary regulator of sex dimorphism in the Drosophila brain (Ito, et al. 1996; Ryner, et al. 1996; Kido and Ito 2002; Demir and Dickson 2005; Manoli, et al. 2005; Stockinger, et al. 2005; Rideout, et al. 2007; Kimura, et al. 2008; von Philipsborn, et al. 2011); and sex dimorphism has been shown to be directed by *fru*. We hypothesized that the conservation in the male-specific Fru^M^ contributes to conservation in male-biased expression. There were 1,771 and 729 genes identified as regulated by *fru* in *D. melanogaster* males and females respectively (Dalton, et al. 2013). Male-biased orthologs were enriched for genes regulated by the Fru^M^ protein in *D. melanogaster* males (χ^2^: *p* < 0.0001) and female-biased orthologs were depleted for signatures of Fru^M^ (χ^2^: *p* = 0.002).

The faster-X hypothesis predicts that genes on the X chromosome evolve faster than those on the autosomes (Haldane 1924b, a; Charlesworth, et al. 1987). When considering the unique properties of the X, in combination with sex-differential effects of alleles, there is increased efficiency of positive selection for X-linked alleles that are recessive and male-beneficial, or dominant and female-beneficial (Wu and Davis 1993; Wu, et al. 1996). In the context of the evolution of sex-biased genes, and in combination with other unique properties of the X, this may result in preferential accumulation of sex-biased genes on the X chromosome over evolutionary time (Rice 1984; Charlesworth, et al. 1987; Oliver and Parisi 2004; Ellegren and Parsch 2007). In comparison, the faster-male theory, a possible explanation of Haldane’s rule (Haldane 1922; Wu and Davis 1993; Turelli and Orr 1995; reviewed in Schilthuizen, et al. 2011), predicts faster evolution of genes related to male reproduction, regardless of location (reviewed in Schilthuizen, et al. 2011). These are not mutually exclusive ideas. We observe an enrichment of genes with sex-biased expression on the X chromosome compared to the autosomes in both species (Supplementary Figure 2A). This is consistent with previous studies in the brain and is predicted by models of sexually antagonistic evolution followed by gain of sex-biased or sex-limited expression (Rice 1984; Khodursky, et al. 2020).

Differences between the X chromosome and autosomes in the evolution of gene expression may be due to changes in regulation of chromatin conformation associated with the X. Consistent with this hypothesis, there were a higher proportions of genes with open chromatin marks detected in both species on the X chromosome compared to the autosomes and conserved male-limited H3K4me3 marks are enriched on the X compared to the autosomes.

Association between chromatin and male- and female-biased expression may be related to evolutionary dynamics between the sexes. Sexual conflicts arise when the optima for a specific trait differ between the sexes and therefore selection differs between the sexes. These conflicts can come in two forms: interlocus and intralocus conflict (reviewed in Rice and Holland 1997; Chapman, et al. 2003; Tregenza, et al. 2006; Bonduriansky and Chenoweth 2009; Cox and Calsbeek 2009; Schenkel, et al. 2018). Intralocus conflict occurs when the optimal fitness of a shared trait/locus is different between males and females, with different alleles favored in males and females. It has been argued that the degree of observed sexual dimorphism can signify the extent to which intralocus sexual conflict has been fully or partially resolved (Cox and Calsbeek 2009). In the whole fly, a small proportion (8.5%) of sex-biased genes have evidence of current sexually antagonistic selection (Innocenti and Morrow 2010), indicating that in the majority of cases, any sex-biased expression observed in this study that is associated with intralocus conflict resolution would be expected to result from a history of partially or fully resolved intralocus conflict, rather than ongoing intralocus conflict. We note that we find no association between the fitness associated genes reported by Innocenti and Morrow (Innocenti and Morrow 2010) and the observed conserved/diverged sex-biased orthologs reported here.

The findings that when female closed chromatin marks are absent in both species the male sex-bias ratio is more similar between the species than when there is a mark in only one species, suggests that the closed chromatin marks may play a role in resolving the ongoing sexual conflict in males. The divergence in the degree of male-bias is associated with female H3K27me2me3 marks which are predicted to reduce expression in females. It is possible that this reflects a mechanism of resolving cases of intralocus conflict in which expression of an allele in females has deleterious effects. However, while male-biased genes on the autosomes show potential suppression of expression in females, female-biased genes (both X and autosomal) in both species lack the association of closed chromatin marks in males. This may suggest that genes with female biased expression either i) don’t involve deleterious effects in males, ii) involve genes that are important for male fitness and are incompatible with closed marks and gene silencing, or iii) do not involve resolution of intralocus sexual conflict. These overall patterns suggest specific testable hypotheses regarding the role of activation and repression via chromatin modifications in the resolution of intralocus sexual conflict for future experiments.

## Methods

### Experimental Design

Isogenic male and female *D. melanogaster* (DGRP r153 and r301) (Mackay, et al. 2012) and *D. simulans* (Winters lines sz11 and sz12) (Signor 2017) flies were raised on standard Bloomington recipe medium at 25C with a 12-h light/dark cycle. There were 2 sexes and 2 genotypes for each species with 6 replicates for a total of 48 samples. Half of the samples were exposed to ethanol. Samples were flash frozen in liquid nitrogen and freeze dried (Supplementary Figure 10).

For RNA-seq, 12 heads from each sample were collected. mRNA purification, cDNA synthesis and dual index barcoding library preparation were carried out by Rapid Genomics (Gainesville, FL, http://rapid-genomics.com). Individual libraries (n=48) were pooled in equimolar ratios as estimated by Qubit and sequenced on a total of 7 Illumina lanes at Rapid Genomics (paired-end 2×100 3 lanes with HiSeq 3000 and paired-end 2×150 2 lanes with HiSeq X and 2 lanes with NovaSeq 6000). External RNA Control Consortium (ERCC) spike-in control was used to evaluate the quality of all RNA-seq sequencing libraries (Jiang, et al. 2011). After the first lane, read counts of each library were used to confirm the pooling strategy.

For ChIP-seq, a target number of ~200 heads from each sample of *D. melanogaster* r301 and *D. simulans* sz11 were collected (2 species x 6 replicates x 2 sexes x 1 genotype = 24 samples). Each sample was used to assay histone marks H3K4me3 (open chromatin), H3K27me2me3 (closed chromatin), and input. (3 antibodies/input x 24 samples = 72 assays). One r301 female untreated sample contained ~175 heads and one 2 sz11 male ethanol treated sample contained ~120 heads, and one sz11 ethanol treated female sample contained 50 heads. A full protocol for the ChIP (Supplementary File 3, developed by NM and RR) is available in Supplementary File 1. ChIP samples were indexed, pooled, and sequenced on one lane of an Illumina HiSeq2500 (paired-end 2×100) at the University of Florida, ICBR (Gainesville, FL, https://biotech.ufl.edu/).

### Genome Annotations

All genome and annotation versions used were from FlyBase release FB2017_04 (http://www.flybase.org) *D. melanogaster* FlyBase r6.17 and *D. simulans* FlyBase r2.02. The FlyBase gene OrthoDB ortholog report (Waterhouse, et al. 2013) (Supplementary File 4) was used to identify one-to-one orthologous gene pairs (one gene in *D. melanogaster* associating with one gene in *D. simulans*, and vice versa).

We created BED files for both genic features (exons, exonic features, TSS +/-150 bp, 5’ UTR, 3’UTR, and introns) and intergenic features (defined as the non-genic features greater than 50 bp in length) for each reference from the relevant GFF annotation file. We note that in areas where there were overlapping exons (where intron/exon boundaries vary by transcript), alternative donor and acceptor sites were defined as exonic and tracked as separate features in downstream analyses (Newman, et al. 2018). Counts of each unique feature type are in Supplementary Table 4. We note that there are fewer genic features annotated in *D. simulans* compared to *D. melanogaster*.

### RNA-seq and ChIP-seq

All results were consistent with reasonable quality data (Yang, et al. 2014) albeit with some shorter sequences and higher duplication rates typically associated with libraries run on the NovaSeq 6000 in some of the RNA-seq runs.

Sequencing adapters were removed from both RNA-seq and ChIP-seq reads using Cutadapt version 2.1 (Martin 2011) with a max error rate of 0.1 and a minimum overlap of 3 nt. Forward and reverse reads were merged using BBMerge (Bushnell, et al. 2017). Reads less than 14bp + 50% original read length were not considered further. Identical reads were identified (fastqSplitDups.py) and removed. The resulting processed reads consisted of i) merged reads (‘single-end’), ii) unmerged reads without a proper pair (‘single-end’), and iii) unmerged reads with proper pairs (paired-end).

Processed RNA-seq reads and all ChIP reads were aligned to the corresponding genome reference (*D. melanogaster* reads mapped to *D. melanogaster* FlyBase r6.17 and *D. simulans* reads mapped to *D. simulans* FlyBase r.202) using BWA-MEM v0.7.15 (Li 2013) as single-end or paired-end with default parameters. To determine if there was any systematic reference bias processed RNA-seq reads from *D. melanogaster* samples were mapped to the *D. simulans* FlyBase r.202 genome, and *D. simulans* samples were mapped to the *D. melanogaster* FlyBase r6.17 genome. A small bias was observed towards mapping to the *D. simulans* genome and in both species, female samples tended to have, on average, slightly higher mapping rates in the ChIP experiment. Sensitivity to mapping bias was examined and results are described in detail in (Supplementary Materials Section 5.3).

### RNA-seq feature detection

A feature was considered detected by RNA-seq if at least one read was present in more than 50% of the replicates for a species-sex combination (e.g., present in at least 7 of the 12 female or male replicates for a given species). The number of detected features for each species-sex combination is summarized in Supplementary Table 5. There are fewer features in *D. simulans* and despite the slightly higher mapping rates found in *D. simulans* samples, there are slightly fewer features detected in *D. simulans* samples compared to *D. melanogaster* across all feature types except for 3’UTR. The 3’UTR features has a higher proportion detection in *D. simulans* compared to *D. melanogaster*, suggesting there may be a systematic bias in the 3’UTR regions of the two species of either an over-annotation of these regions in *D. melanogaster* or an under-annotation in *D. simulans*. There do not seem to be many missing genes in the *D. simulans* annotation because there are not more detected features in *D. simulans* intronic and intergenic features compared to *D. melanogaster*. In fact, there is a lower proportion of detected intronic and intergenic features in *D. simulans* samples compared to *D. melanogaster* samples. Exonic feature detection was similar between the species, with a slightly higher detection rates in *D. melanogaster* males. A feature was considered sex-limited if the feature was detected in only one of the 2 sexes. Approximately 4% of exonic features were sex-limited in *D. melanogaster* samples (2,530 in males, 1,195 in females) and *D. simulans* (1,801 in males, 1,506 in females).

For the gene expression analysis, exonic regions were separated into non-overlapping exonic features where alternative donor/acceptor sites were quantified separately from shared exonic regions, in order to capture the potential sex-specific structures in the gene (Newman, et al. 2018). Genes were defined as detected if at least one exonic feature was detected for either sex. There are a similar number but proportionally more genes detected in *D. simulans* (11,543 out of 15,385, ~75%) compared to *D. melanogaster* (11,716 out of 17,737, ~66%) indicating that there were no large quality differences in the *D. simulans* genome compared to the *D. melanogaster* samples to the *D. melanogaster* genome despite the differences in annotation.

To compare genes across *D. melanogaster* and *D. simulans*, we focus on annotated orthologs from the OrthoDB ortholog report (Waterhouse, et al. 2013) to identify one-to-one orthologous gene pairs (one gene in *D. melanogaster* associating with one gene in *D. simulans*, and vice versa) (Supplementary File 4). There are 14,241 orthologous gene pairs between the species, 12,083 of which are one-to-one orthologs. Genes on chromosome 4, the Y chromosome, and scaffolds of either species were excluded from further analysis. There were 7 genes on the X chromosome of *D. melanogaster* with orthologs on autosomes of *D. simulans*, and 1 gene on the X of *D. simulans* with an ortholog on an autosome of *D. melanogaster*. These 8 genes were also excluded. The remaining 11,937 one-to-one orthologous genes on the X (n = 1,840) and autosomes (n = 10,097) of both species were carried forward.

### RNA-seq Differential Expression

For each species, exonic features were quantified as *C_is_* = (∑(*d_ijs_*)/*N_i_*) × (*Q*/*U_s_*), where *d* is the depth of reads at nucleotide *j* of feature *i*, *N* is the length of the feature, *U_s_* is the upper quartile of (∑(*d_ijs_*)/*N_i_*) values in sample s, and *Q* is the median of all *U_s_* values within the given species (Bullard, et al. 2010; Dillies, et al. 2013) (Supplementary File 5). Distributions of upper quartile values across exonic features were evaluated for each sample mapped to the genome of the sample species (Supplementary Figure 11). Median upper quartile values and associated distributions were strikingly similar across all samples in both species except for one *D. simulans* sz12 male replicate, which was removed from further analysis.

For each species separately, differential expression between males and females was evaluated for exonic features detected in both sexes. We used the linear fixed effect model *Y_xp_* = *μ* + *g_x_* + *ε_xp_*, where *Y* is the log-transformed UQ normalized *C_is_* values for the *x*th sex (*x* = male, female), *p*th replicate (*p* = 1,2,…,12). We accounted for potential heteroscedasticity of variance between the sexes (Graze, et al. 2012) and used the Kenward-Roger adjustment for the degrees of freedom (Kenward and Roger 1997). Normality of residuals was tested using the Shapiro-Wilk test (Shapiro and Wilk 1965). Fold-change ratios were calculated for each exonic feature *i*, *r_i_* = (∑(*f_ip_*)/*k*)/(∑(*m_il_*)/*n*), where *f_ij_* is the UQ normalized *C_is_* for exonic region *i* in female replicate *p* = 1…*k* total female replicates, and *m_il_* is the UQ normalized *C_is_* for exonic region *i* in male replicate *l* = 1…*n* total male replicates. Exonic features were classified as male-biased (or female-biased) if the nominal p-value was less than or equal to 0.05 and the fold-change less than (or greater than) 1.

### ChIP-seq Feature Detection

While peak calling is a common method of ChIP-seq analysis; it is highly dependent on the algorithm used and the parameters selected (Yang, et al. 2014), especially for ChIP marks that are predicted to show broad peaks such as certain histone modifications (Park 2009; Pepke, et al. 2009; Dahl, et al. 2016). To have a consistent method for evaluation and comparison of ChIP results across different marks and between males and females, and to compare ChIP results directly to the RNA-sea results in *cis*, we use ChIP-seq reads to quantified features based on the annotations of the reference genomes (Katz, et al. 2010; Anders, et al. 2012; Zhang, et al. 2012; Yang, et al. 2014; Newman, et al. 2018). By focusing on features rather than MACS2 peaks, many more detections above input control are identified at the feature-level and at the gene-level (See Supplementary Methods Section 7.1 for detailed results from MACS2).

A feature was considered detected above the input control in H3K4me3/H3K27me3me4 (DAI) if *C*_*K*4,is_ > *C_Input,is_*, in more than 50% of the replicates for that species-sex combination A gene was considered as having a mark if at least one exonic feature in the gene was DAI. A gene was considered male-limited (or female-limited) if only sex-limited exonic features were identified in both treatments. The agreement between histone marks for males and females, as well as between H3K4me3 and H3K27me2me3 marks within each sex, was estimated using Cohen’s kappa (Fleiss 1981) rather than simple agreement in order to account for marginal frequencies and provide a more accurate assessment of the relationship between sexes and the marks (Supplementary Figure 4).

### Chromatin and expression

Histone modifications change the availability of chromatin for transcription (Santos-Rosa, et al. 2002; Schneider, et al. 2004; Wang, et al. 2008; Juan, et al. 2016); therefore, we examine the impact of chromatin marks on expression. When sex-biased expression is observed, this may be due to open marks in the sex with the higher expression, or closed marks in the other sex. Specifically, if there is male-biased expression, are there open (H3K4me3) marks in males or closed (H3K27me2me3) marks in females for that gene, and if there is female-biased expression, are there open (H3K4me3) marks in females or closed (H3K27me2me3) marks in males (Figure 3). As chromatin marks in males do not influence expression in females, or vice versa, the appropriate statistical comparison is not a test of general association between expression and chromatin marks between the sexes.

For males, the presence/absence of the chromatin marks, H3K4me3 and H3K27me2me3, was compared to presence/absence of gene expression in males and evaluated for agreement using Cohen’s kappa coefficients (Fleiss 1981) (Supplementary Table 3). Females were examined separately in the same manner. For genes with detected expression in both sexes, the presence/absence of sex bias in males was compared to the presence/absence of male H3K4me3 marks using Fisher exact test (Fisher 1934) with the alternative expectation that male open chromatin marks would be more likely in male-biased expression. For genes with sex-biased expression in males, the presence/absence of H3K27me2me3 marks in females was tested using Fisher exact test (Fisher 1934) with the alternative expectation that female closed chromatin marks would be more likely in genes with male-biased expression. Tests were similarly performed for the presence/absence of sex bias in females compared to the presence/absence of female H3K4me3 and presence/absence of male H3K27me2me3 using Fisher exact test (Fisher 1934).

### List enrichment

Genes with sex-biased gene expression conserved between *D. melanogaster* and *D. simulans* in this study were compared to genes identified in previous studies of sex-biased expression in *D. melanogaster* head tissue (Chang, et al. 2011) using Pearson’s Chi-square (χ^2^) test (Pearson 1900). Additionally, conserved male-biased (or female-biased) genes were compared to genes previously identified as male-biased (or female-biased) in *D. melanogaster* head tissue and in *fru-P1*-expressing neurons (Newell, et al. 2016) using Pearson’s Chi-square (χ^2^) test (Pearson 1900). Based on the extensive knowledge of the sex-specifically spliced Drosophila sex determination gene *fru* (Ryner, et al. 1996; Heinrichs, et al. 1998; reviewed in Salvemini, et al. 2010), we expected *fru* to play a role in conserved sex-biased expression. Genes with male-biased and female-biased expression conserved between *D. melanogaster* and *D. simulans* in this study were compared to genes regulated by the Fru^M^ protein in *D. melanogaster* males (Dalton, et al. 2013) using Pearson’s Chi-square (χ^2^) test (Pearson 1900).

Divergence of the targets of the terminal sex determination genes may contribute to the divergence of sex-biased expression between the species. To evaluate this, species-specific sex-biased genes identified in this study were compared to a genes in a study of *dsx* regulation in *dsx* null females and *dsx* pseudomales of *D. melanogaster* (Arbeitman, et al. 2016) and to genes observed to be regulated downstream of *fru* in *D. melanogaster* males (Dalton, et al. 2013) using Pearson’s Chi-square (χ^2^) test (Pearson 1900). To validate the patterns of open and closed chromatin in males and females, gene-level presence of open (H3K4me3) and closed (H3K27me2me3) chromatin marks in *D. melanogaster* males and females found in this study were compared to previous observations of H3K4me3 and H3K27me3 marks in *D. melanogaster* male and female (*elav*-expressing) neurons (Palmateer, et al. 2021) using Pearson’s Chi-square (χ^2^) test (Pearson 1900). Tests of agreement between these datasets were carried out for males and females separately using Cohen’s kappa coefficients (Fleiss 1981) (Supplementary Table 3).

To evaluate if the patterns of the chromatin marks in the head tissue described here are consistent with patterns of chromatin marks in neurons known to direct male and female reproductive behaviors (Demir and Dickson 2005; Manoli, et al. 2005; Stockinger, et al. 2005; Kvitsiani and Dickson 2006), the genes we detected with open (or closed) chromatin marks were compared to genes with H3K4me3 (or H3K27me3) marks in *D. melanogaster* male and female *fru-P1-* expressing neurons (Palmateer, et al. 2021) using Pearson’s Chi-square (χ^2^) test (Pearson 1900). We also compared genes with male-limited and female-limited open (or closed) chromatin to the genes with H3K4me3 (or H3K27me3) marks in *D. melanogaster* male and female *fru-P1-* expressing neurons (Palmateer, et al. 2021) using Pearson’s Chi-square (χ^2^) test (Pearson 1900). Tests of agreement of the comparable marks between head tissue and *fru-* P1-expressing neurons were also evaluated for males and females separately using Cohen’s kappa coefficients (Fleiss 1981) (Supplementary Table 3).

## Data availability

Raw short-read data from the RNA-seq and ChIP-seq experiments are available under SRA BioProject accession PRJNA737411. RNA-seq and ChIP-seq mapped read count summary (Supplementary Table 6) and RNA-seq UQ normalization factors (Supplementary File 5) are provided in the supplement. Analyzed data are provided as supplementary files for i) *D. melanogaster* gene-level chromatin and expression variables (Supplementary File 1), ii) *D. melanogaster* feature-level level chromatin and expression variables (Supplementary File 6), iii) *D. simulans* gene-level chromatin and expression variables (Supplementary File 2), iv) *D. simulans* feature-level chromatin and expression variables (Supplementary File 7), and v) *D. melanogaster-D. simulans* orthologous gene chromatin and expression variables (Supplementary File 8). Further detail of methods can be found in Supplementary Materials and documentation of all analyses and comparisons as well as scripts are on github (https://github.com/McIntyre-Lab/papers/tree/master/nanni_chip_rna_2022).

## Acknowledgments

Michelle Arbeitman for helpful discussion and sharing of pre-publication results. HiPerGator high-performance supercomputer at the University of Florida. The University of Florida Genetics Institute. Funding provided by National Institute of General Medical Sciences R01GM128193 (LMM, SVN) and National Institute of Dental and Craniofacial Research R01DE026707 (RR).

## Supplementary Figures

**Supplementary Figure 1.**
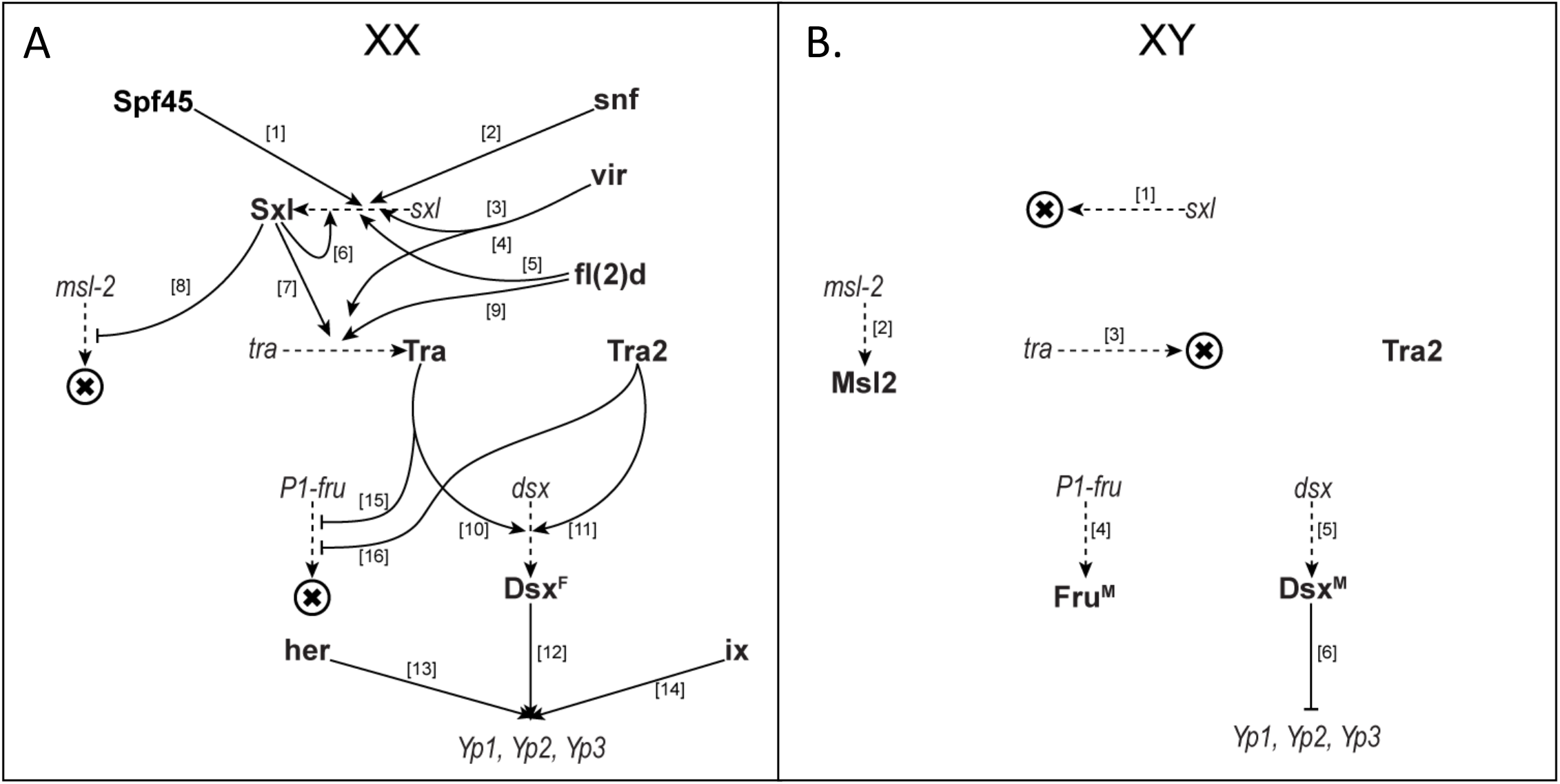
Drosophila sex determination hierarchy, XX females (A) and XY males (B) adapted from Figure 1 in Fear, et al. 2015. Transcripts are italicized and proteins are bold. Solid arrows are genetic interactions (e.g., splicing, transcription) and dashed arrows are protein translation. The X within a circle represents no productive protein product. (A1) Spf45 → Sxl (Lallena, et al. 2002), (A2) Snf → Sxl (Flickinger and Salz 1994), (A3) vir→ Sxl (Hilfiker, et al. 1995), (A4) vir → tra (Hilfiker, et al. 1995), (A5) fl(2)d → Sxl (Granadino, et al. 1990), (A6) Sxl → Sxl (Cline 1978; Bell, et al. 1988; Lallena, et al. 2002), (A7) Sxl → Tra (Sosnowski, et al. 1989; Inoue, et al. 1990), (A8) Sxl H Msl-2 (Bashaw and Baker 1997; Kelley, et al. 1997; Gebauer, et al. 1998), (9A) fl(2)d → Tra (Granadino, et al. 1996), (A10) Tra → Dsx^F^ (Inoue, et al. 1992), (A11) Tra2 → Dsx^F^ (Inoue, et al. 1992), (A12) Dsx^F^ → Yps (Burtis, et al. 1991; Coschigano and Wensink 1993; An and Wensink 1995; Erdman, et al. 1996), (A13) Her → Yps (Li and Baker 1998), (A14) ix→Yps (Garrett-Engele, et al. 2002), (A15) Tra ┤ Fru^M^ (Ryner, et al. 1996; Heinrichs, et al. 1998), (A16) Tra2 ┤ HFruM (Ryner, et al. 1996; Heinrichs, et al. 1998), (B1) default splicing of *sxl* transcripts results in no functional protein (Bell, et al. 1988), (B2) Msl-2 protein produced (Bashaw and Baker 1995; Kelley, et al. 1995; Zhou, et al. 1995), (B3) default splicing of *tra* transcripts results in no functional protein (Boggs, et al. 1987), (B4) Fru^M^ protein produced (Ryner, et al. 1996; Heinrichs, et al. 1998), (B5) default splicing of *dsx* transcripts in XY individuals results in Dsx^M^ protein (Burtis and Baker 1989), (B5) Dsx^M^ represses expression of Yps (Coschigano and Wensink 1993).

**Supplementary Figure 2.**
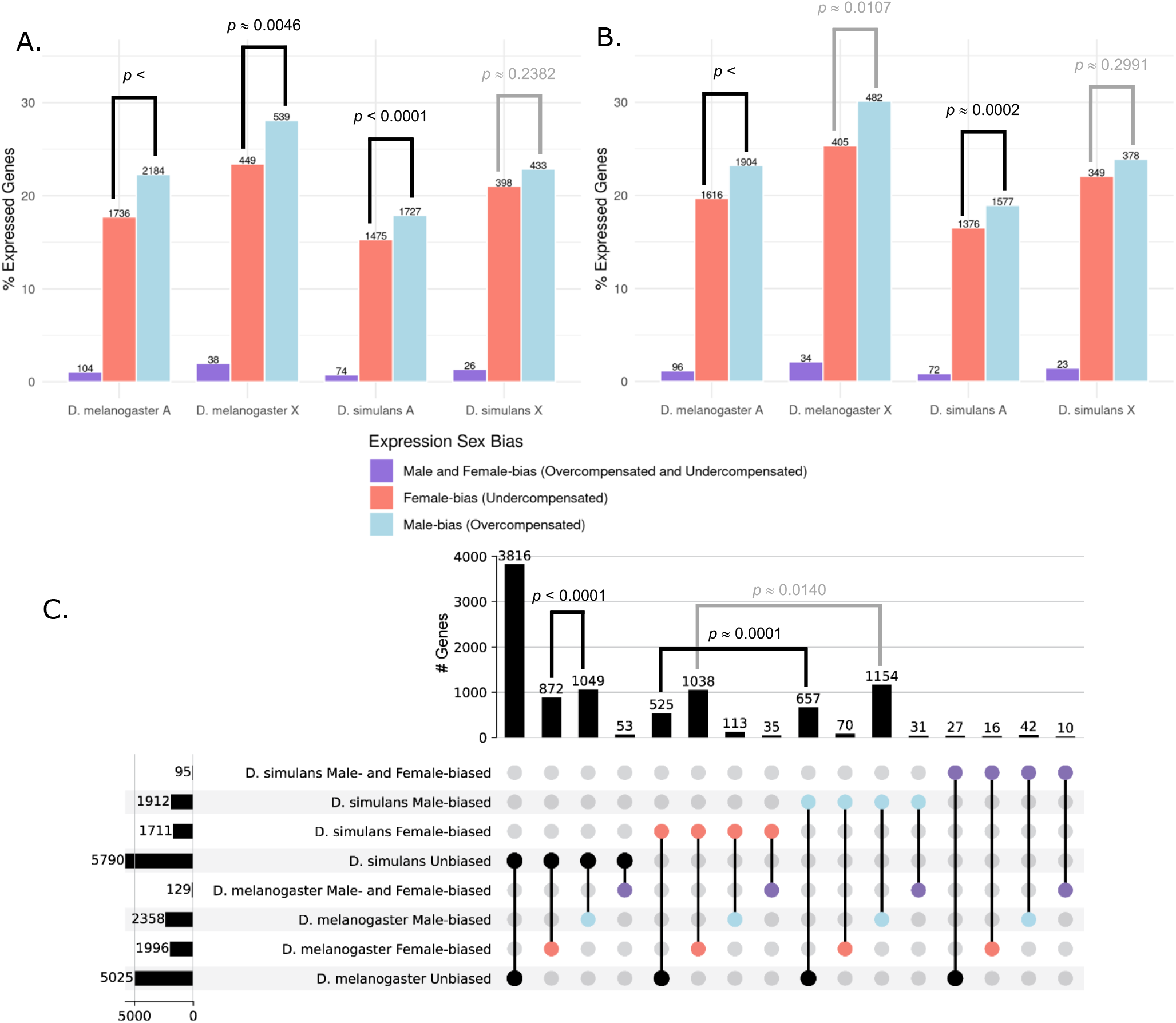
Excess of male-bias in *D. melanogaster* and *D. simulans* is due to divergent male-biased expression. (Panel A) Relative percent of expressed genes for *D. melanogaster* on the X (X, 1,919) and autosomes (A, 9,797) and for *D. simulans* on the X (X, 1,893) and autosomes (A, 9,650). (Panel B) Relative percent of orthologous genes expressed in *D. melanogaster* X (1,599) and autosomes (8,206) and *D. simulans* X (1,583) and autosomes (8,327). Chromosome 4 is excluded from the autosomes. Note that total numbers of orthologous genes differ between the species due to differences in chromosomal assignments. The number of genes in each category is printed over the box with male-bias (blue), female-bias (red), and both male- and female-biased (purple). Sex-limited genes (expressed in only one sex) are excluded. P-values for differences in male-biased and female-biased expression are reported. (Panel C) Conservation and divergence of sex-biased expression for expressed orthologous genes (n=9,508, with a consistent X/autosome chromosomal assignment between the species). Dots below the histogram are solid for the combination of factors reported as the number of genes in the bar plot.

**Supplementary Figure 3.**
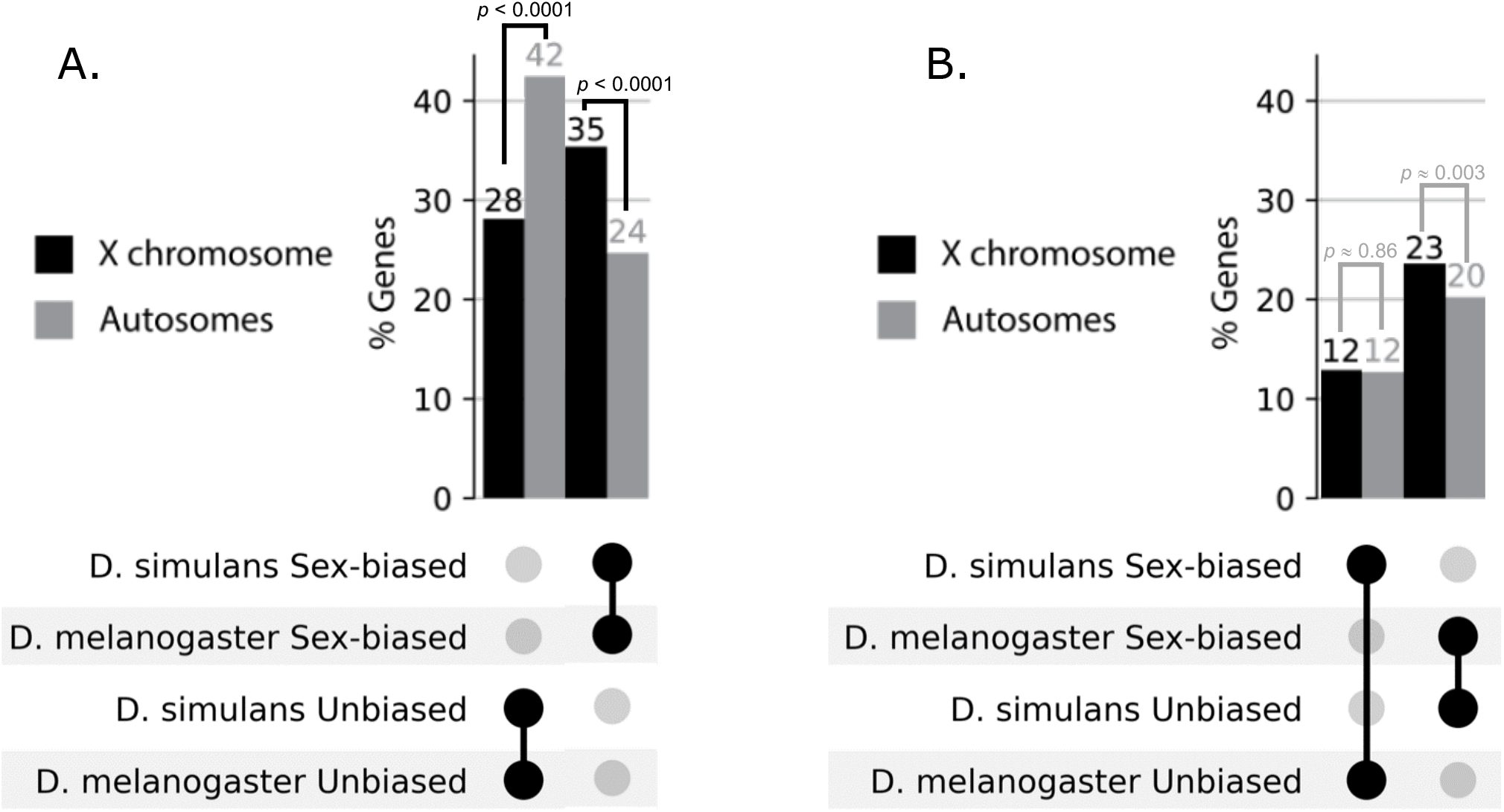
X vs. autosomes of orthologs with conserved and divergent sex-biased expression. Expression of orthologous genes in the head for both sexes and both species on the X (n_x_=1,529 genes) and autosomes (n_a_= 7,979). (Panel A) Genes that are conserved in their sex bias (n_x_=541, n_a_= 1,968) are more likely to be on the X (35% on X vs. 24% on autosomes; χ^2^: *p* < 0.0001), while those that are unbiased are more likely to be on the autosomes (28% on X vs. 42% on autosomes; χ^2^: *p* < 0.0001). (Panel B) Genes divergent in sex bias (n_x_=558, n_a_= 2,625) have no significant chromosomal bias for either *D. simulans*-specific sex-biased genes (12% on X vs. 12% on autosomes; χ^2^: *p* = 0.86) or *D. melanogaster-specific* sex-biased genes (23% on X vs. 20% on autosomes; χ^2^: *p* = 0.003). Connected black dots indicate the category plotted in the two bars above. The Y-axis is of the percentage of the total number of genes on the X or autosomes in each of the four categories. Chromosome 4 is excluded from the autosomes. X vs. autosome tests were performed using Pearson’s Chi-square (χ^2^) test (Pearson 1900) with a significance threshold of *p* < 0.001.

**Supplementary Figure 4.**
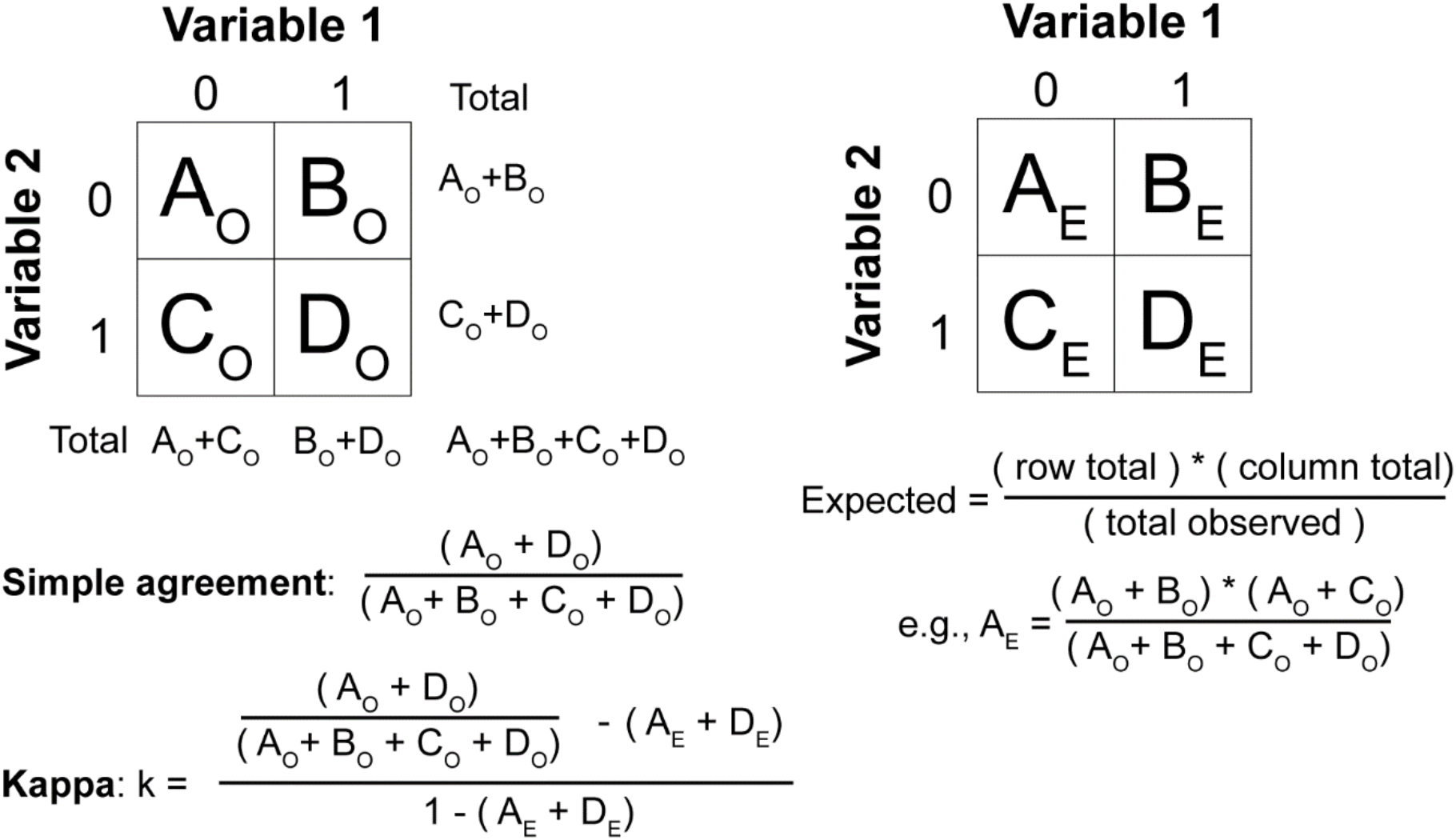
Measurements of Agreement. Given the table of observed values A_O_, B_O_, C_O_, and D_O_, the expected values indicated on the right can be calculated (A_E_, B_E_, C_E_, and D_E_). The formulas for calculating simple agreement and Cohen’s Kappa agreement (Fleiss 1981) are also presented. Cohen’s Kappa values correct for marginal frequencies, for when there is an imbalance between the variables tested.

**Supplementary Figure 5.**
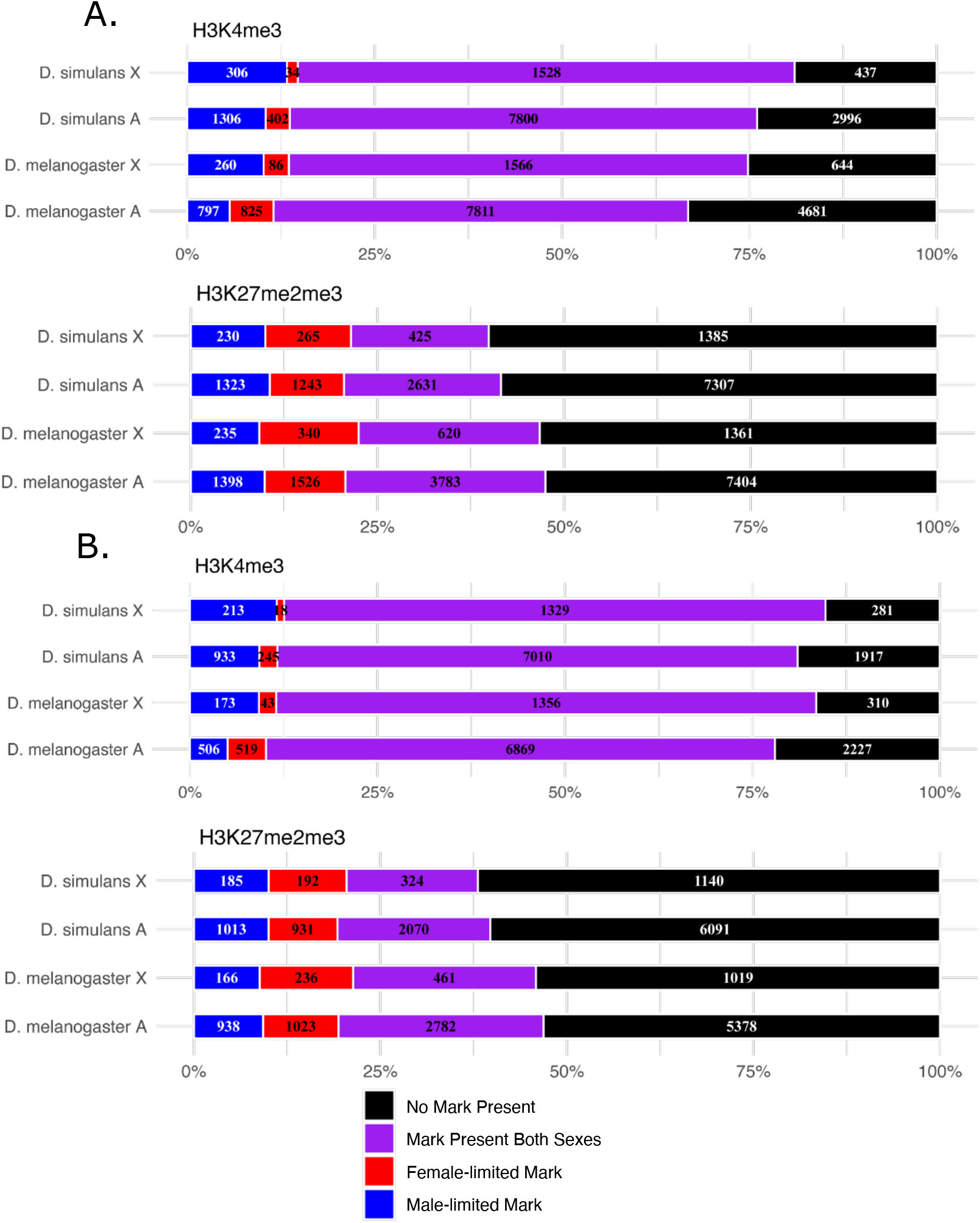
Chromatin marks in males and females. (Panel A) Genes on the X or autosomes (denoted as A) of *D. melanogaster* FlyBase reference r6.17 (n_x_=2,556; n_A_=14,114; n_x_ + n_A_ = 16,670) and *D. simulans* FlyBase reference r2.02 (n_x_=2,305; n_A_=12,504; n_x_ + n_A_ =14,809) with the number of genes with H3K4me3 (top) or H3K27me2me3 (bottom) male-limited, female-limited, or detected in both sexes indicated in blue, red, and purple respectively. Note that chromosome 4 is not included in the autosomes. (Panel B) Similar to Panel A, but with selecting for the one-to-one orthologs between *D. melanogaster* and *D. simulans* (n=12,083), excluding genes on chromosome 4 or unmapped scaffolds from further analysis (80 genes in *D. melanogaster* and 137 genes in *D. simulans*), resulting in 1,882 and 10,121 genes are on the *D. melanogaster* X and autosomes respectively, and 1,841 and 10,105 for *D. simulans* X and autosomes.

**Supplementary Figure 6.**
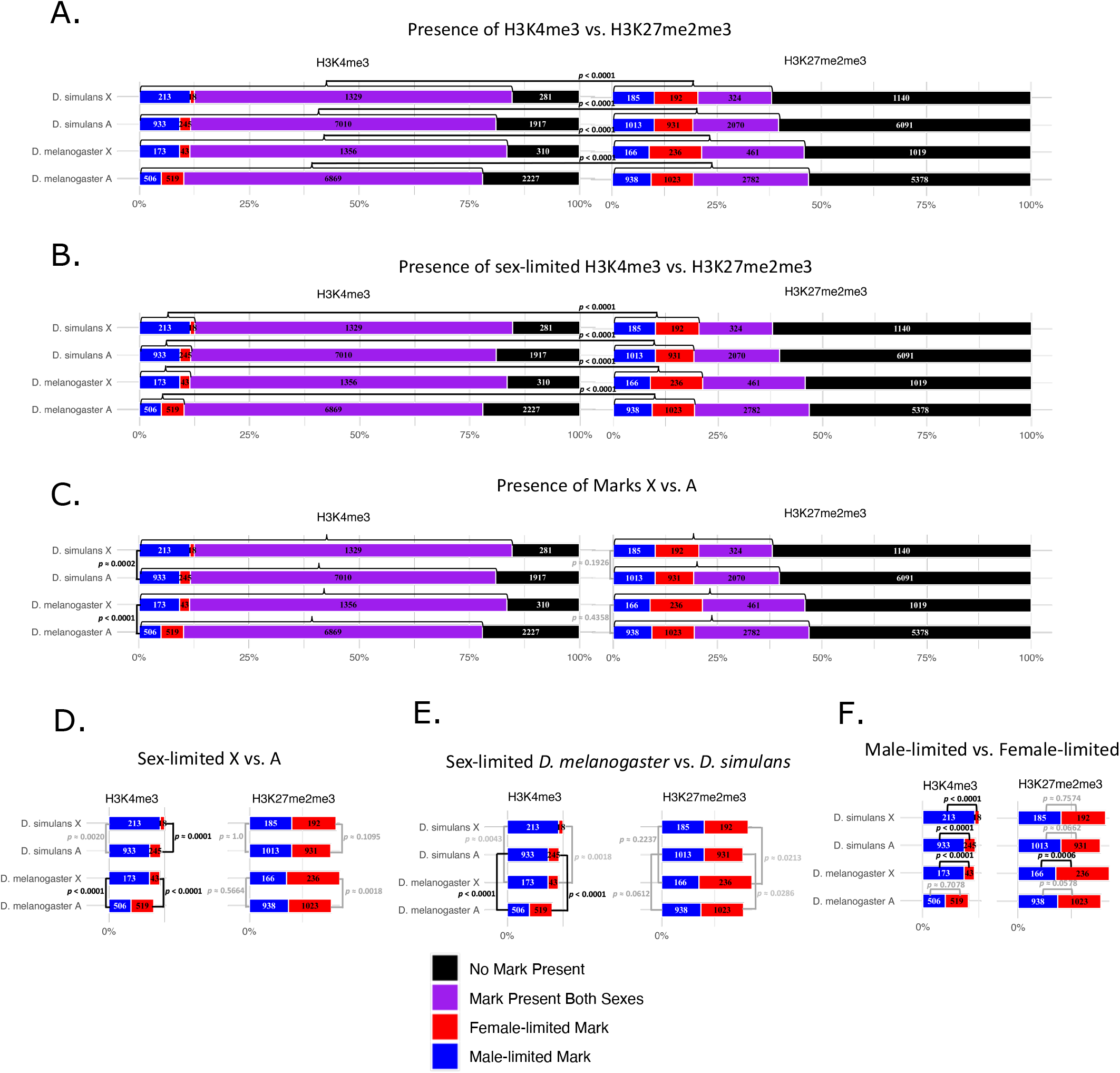
Tests for differential chromatin. H3K4me3 and H3K27me2me3 marks not present (black), present only in females (female-limited, red), present only in males (male-limited, blue), or present in both males and females (purple) on the X chromosome (X) and autosomes (A) for one-to-one orthologous genes of *D. melanogaster* and *D. simulans*. Chromosome 4 is excluded from the autosomes. The number of genes for each group is indicated. The total genes evaluated for are 1,882 and 10,121 for *D. melanogaster* X and autosomes respectively, and 1,841 and 10,105 for *D. simulans* X and autosomes. Tests are performed as follows. Panel A compares the presence of H3K4me3 vs. H3K27me2me3 marks in males/females within each species and chromosomal location. Panel B compares the presence of sex-limited H3K4me3 vs. H3K27me2me3 marks within each species and chromosomal location. Panels C-F compare within H3K4me3 or H3K27me2me3 marks separately. Panel C compares the presence of chromatin marks in either sex on the X vs. the autosomes within each species.Panel D compares the presence of male-limited or female-limited marks on the X vs. the autosomes within each species. Panel E compares the proportion of male-limited or female-limited marks between D. melanogaster and D. simulans within each chromosomal location. Panel F compares male-limited vs. female-limited within each species and chromosomal location. All tests of X vs. autosomes are evaluated using Pearson’s Chi-square (χ^2^) test (Pearson 1900). Differences between species, sexes, and H3K4me3 vs. H3K27me2me3 are evaluated using McNemar’s test of homogeneity (McNemar 1947).

**Supplementary Figure 7.**
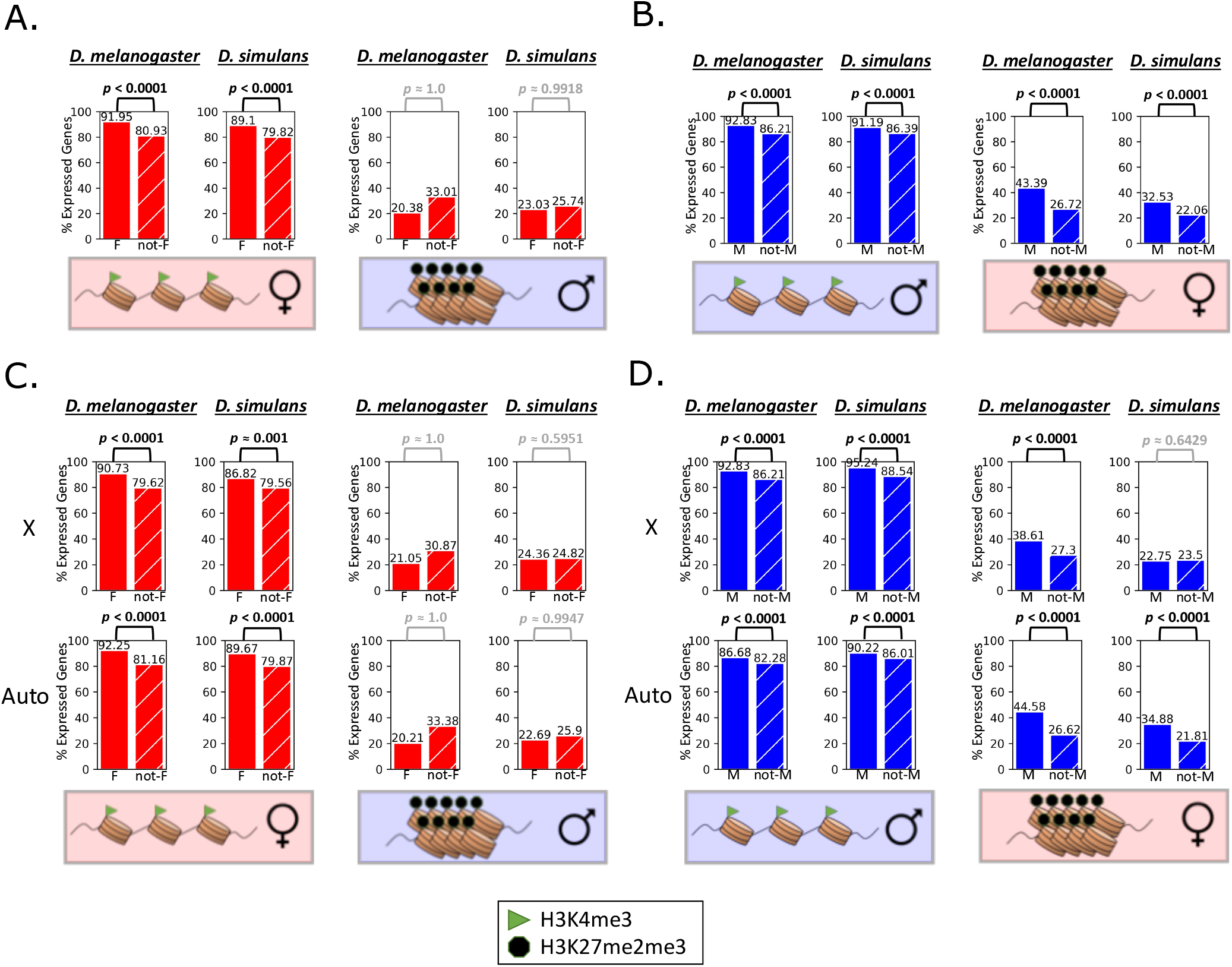
Sex-biased expression is associated with chromatin marks in subset of orthologs. The Y-axis of each graph represents the percent of expressed female-biased (solid red), nonfemale-biased (hatched red), male-biased (solid blue), or non-male-biased (hatched blue) genes with a one-to-one ortholog within each species with the indicated chromatin (cartoon representations below each set of bars). Consistent with the model presented in Figure 4, (Panel A) Female-biased genes (solid red) are enriched for H3K4me3 (open) chromatin when compared to non-female-biased genes (hatched red) in both species. (Panel B) Male-biased genes (solid blue) are enriched for male open chromatin and female H3K27me2me3 (closed) chromatin when compared to non-male-biased genes (hatched blue) in both species. The model in Figure 4 was also evaluated for X and autosomes separately. (Panel C) Female-biased genes (solid red) are enriched for open chromatin when compared to non-female-biased genes (hatched red) on both the X and autosomes of both species. (Panel D) Male-biased genes (solid blue) are enriched for male open chromatin and female closed chromatin when compared to non-male-biased genes (hatched blue) on both the X and autosomes of *D melanogaster. D simulans* shows the same pattern on the autosomes. On the X chromosome, male-bias genes are enriched for open chromatin in males but not for closed chromatin in females, showing a divergence in the regulatory pattern between the two species. There were 11,937 orthologous genes evaluated, 9,747 (n_x_=1,562, n_A_=8,182) genes expressed in *D. melanogaster* and Y genes expressed in *D. simulans* (n_x_=1,582, n_A_=8,320). Each set of female-biased (male-biased) and non-female-biased (non-male-biased) genes were tested for enrichment of the indicated chromatin mark using Fisher exact test (Fisher 1934) with the alternative expectation that the indicated chromatin marks would be more likely in genes with female-biased (male-biased) expression. Significant p-values (*p* < 0.001) are black and p-values above the significance threshold are gray.

**Supplementary Figure 8.**
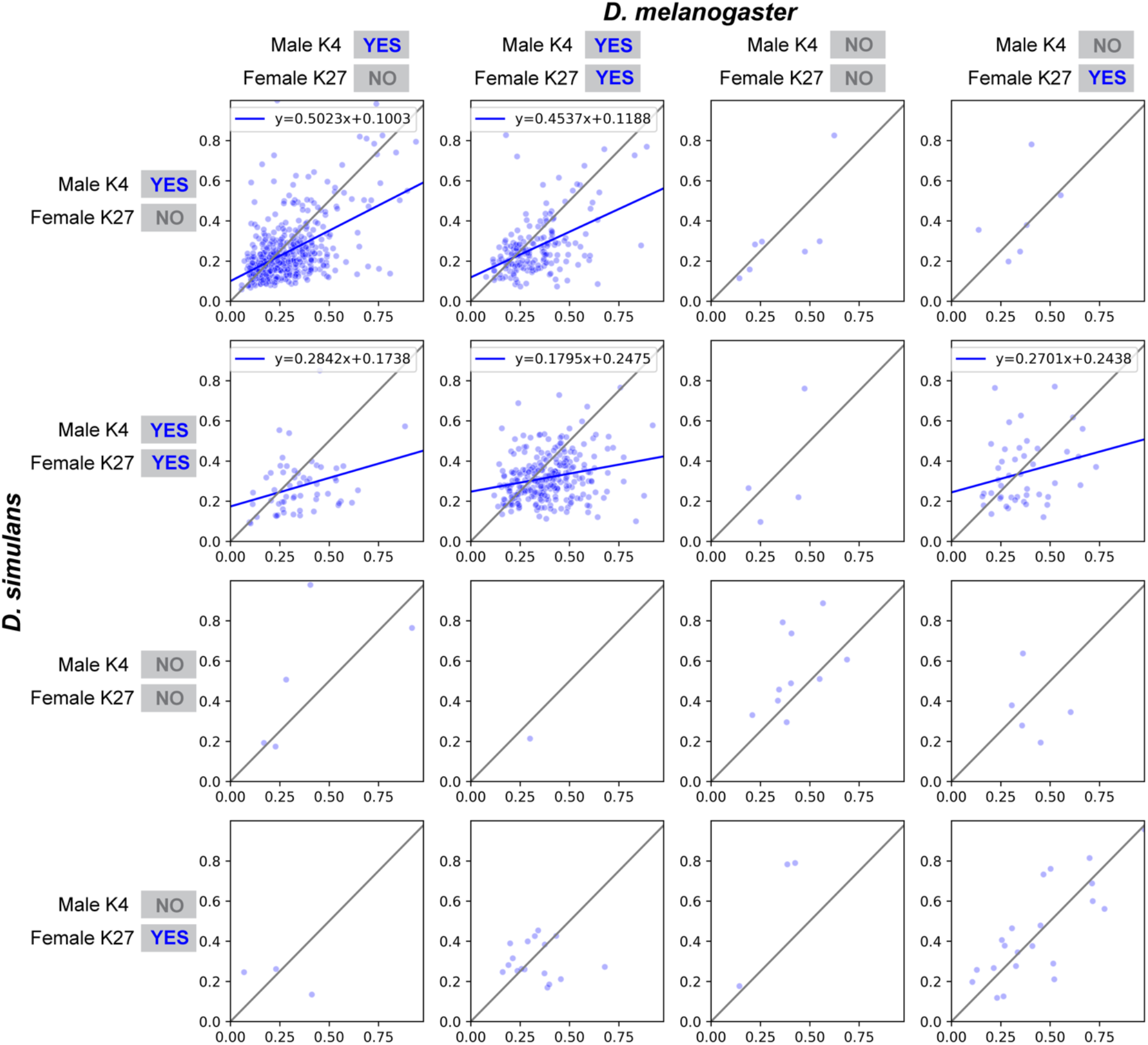
Plotted on the interval of (0,1) is the value 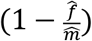 for male-biased orthologs (blue dots), where 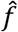 is average UQ normalized expression across female samples and 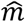 is average UQ normalized expression across male samples. *D. melanogaster* is on the X-axis and *D. simulans* on the Y axis. Each male-biased ortholog is plotted based on the sex bias ratio observed in each species, and placed in the box corresponding to the chromatin observed in *D. melanogaster* and *D. simulans*. Chromatin of *D. melanogaster* is indicated at the top of each column of plots and chromatin of *D. simulans* is indicated at the left of each row of plots. Plots along the diagonal from the top left to the bottom right are genes where the observed chromatin is the same between the species. For each row (*D. simulans*) and column (*D. melanogaster*) the presence of H3K4me3 in males is indicated by a blue “YES” next to “Male K4” or a gray “NO” if it is not present. Similarly for the presence of H3K27me2me3 in females indicated by a blue “YES” next to “Female K27” if present, or a gray “NO” otherwise. Linear regression estimates are calculated for plots with at least 25 genes and plotted as a blue line.

**Supplementary Figure 9.**
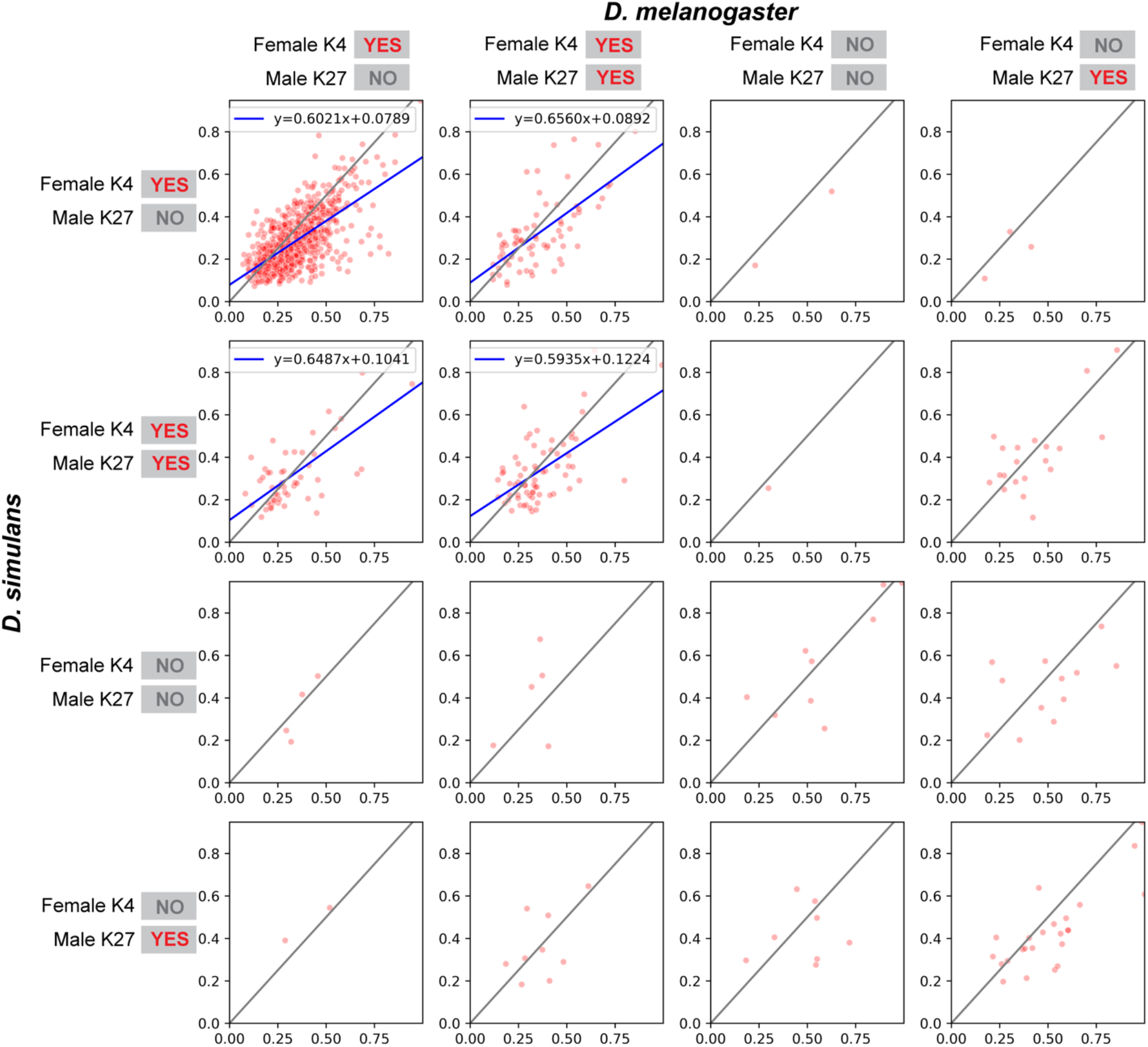
Plotted on the interval of (0,1) is the value 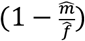 for female-biased orthologs (red dots), where 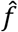 is average UQ normalized expression across female samples and 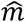 is average UQ normalized expression across male samples. *D. melanogaster* is on the X-axis and *D. simulans* on the Y axis. Each female-biased ortholog is plotted based on the sex bias ratio observed in each species, and placed in the box corresponding to the chromatin observed in *D. melanogaster* and *D. simulans*. Chromatin of *D. melanogaster* is indicated at the top of each column of plots and chromatin of *D. simulans* is indicated at the left of each row of plots. Plots along the diagonal from the top left to the bottom right are genes where the observed chromatin is the same between the species. For each row (*D. simulans*) and column (*D. melanogaster*) the presence of H3K4me3 in females is indicated by a red “YES” next to “Female K4” or a gray “NO” if it is not present. Similarly for the presence of H3K27me2me3 in males indicated by a red “YES” next to “Male K27” if present, or a gray “NO” otherwise. Linear regression estimates are calculated for plots with at least 25 genes and plotted as a blue line.

**Supplementary Figure 10.**
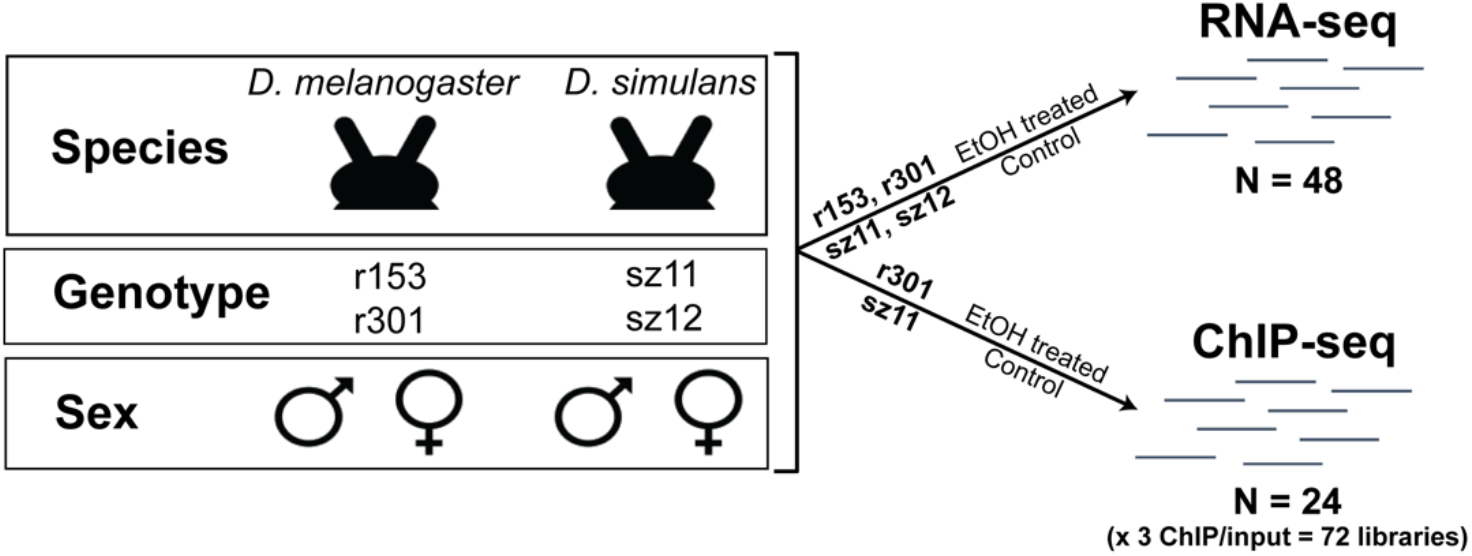
Experimental Design. For RNA-seq there were a total of 48 samples (2 species x 2 genotypes x 2 sexes x 6 replicates). For ChIP-seq there were a total of 24 samples (2 species x 1 genotype x 2 sexes x 6 replicates) used for assaying chromatin (3 antibody/inputs per sample). Note that half of the replicates were exposed to ethanol (EtOH) and are included as additional data.

**Supplementary Figure 11.**
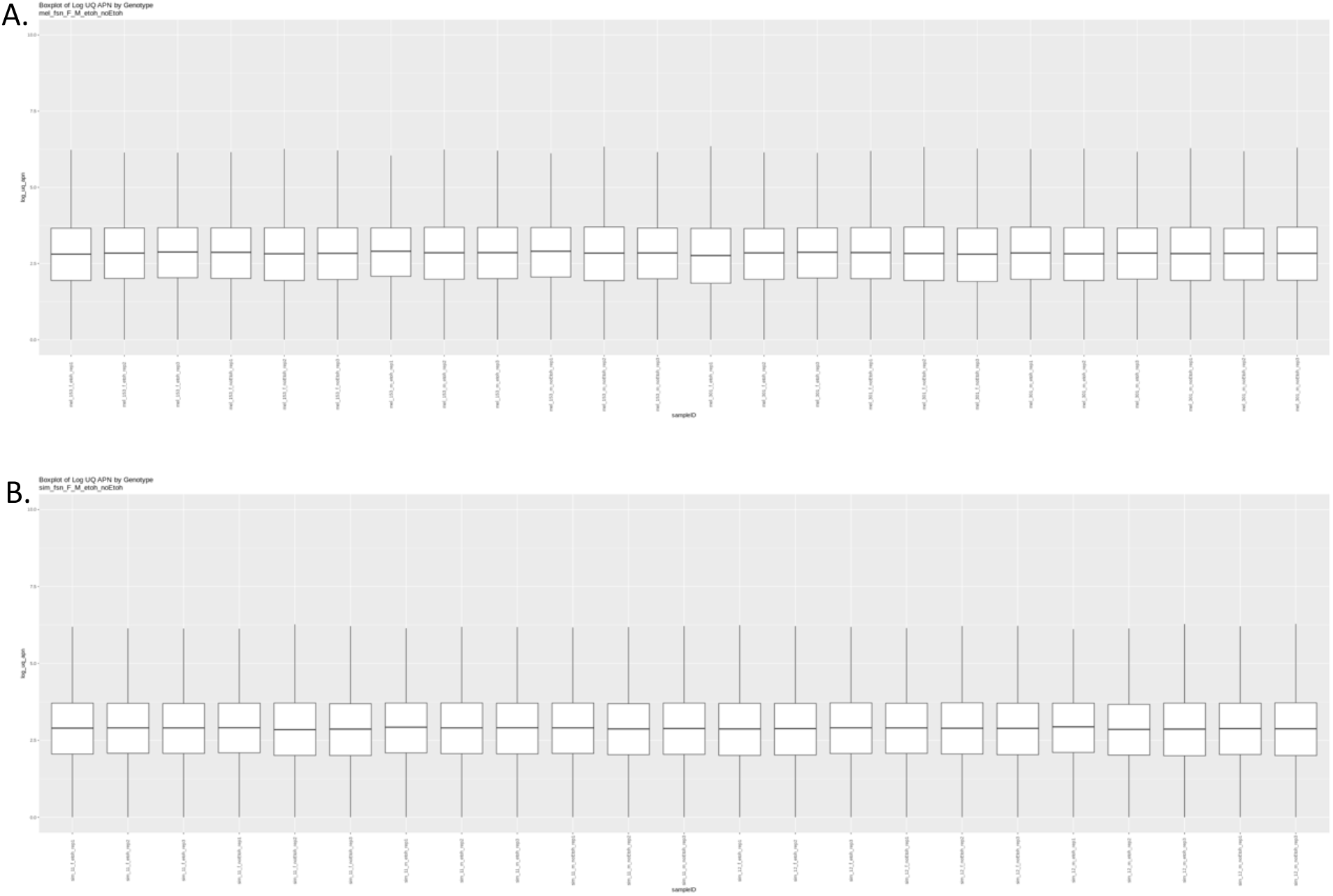
Distributions of RNA-seq expression values after UQ normalization. Upper quartile (UQ) normalization distributions per sample for (Panel A) *D. melanogaster* and (Panel B) *D. simulans* samples excluding the *D. simulans* sz12 male replicate that was removed due to a low median UQ relative to the rest of the samples.

## Supplementary Tables

**Supplementary Table 1.** Number of genes showing different patterns of expression bias. The number of genes on the X and autosomes (excluding chromosome 4) for each pattern of expression bias for *D. melanogaster* and *D. simulans* head tissue (individual counts of X and autosomes are in parentheses). Sex-biased genes are the sum of male-biased (Male), female-biased (Female), and male- and female-biased (Male and Female) genes. Expression bias of orthologous of the species are indicated in the right-most column. Conserved expression bias, where both species are classified as the same category within the orthologous gene pair, are included in rows 1-5, followed by rows 6-18 with diverged expression bias, where the gene pair is assigned different expression categories between *D. melanogaster* and *D. simulans*. Binomial test probabilities are indicated to the right of the table for the comparison of male-biased vs. female-biased for conserved and species-specific sex-biased genes. Significant p-values are in black if below the significant threshold of *p* = 0.001 and gray if above the threshold.

|  |  | *D. melanogaster*<br>(X, A) | *D. simulans*<br>(X, A) | Orthologs<br>(X, A) |  |
| --- | --- | --- | --- | --- | --- |
| 1 | Male-biased | 2723 (539, 2184) | 2160 (433, 1727) | 1154 (235, 919) | } $p = 0.014$ |
| 2 | Female-biased | 2185 (449, 1736) | 1873 (398, 1475) | 1038 (215, 823) |  |
| 3 | Male- and Female-biased | 142 (38, 104) | 100 (26, 74) | 10 (2, 8) |  |
| 4 | Sex-biased | 5050 (1026, 4024) | 4133 (857, 3276) | 2202 (452, 1750) |  |
| 5 | Unbiased | 6666 (893, 5773) | 7410 (1036, 6374) | 3816 (430, 3386) |  |
| 6 | Switch |  |  | 3490 (647, 2843) |  |
| 7 | Reversal | Male | Female | 113 (37, 76) |  |
| 8 | Reversal | Female | Male | 70 (17, 53) |  |
| 9 | Gain/Loss | Male | Male and Female | 42 (12, 30) |  |
| 10 | Gain/Loss | Female | Male and Female | 16 (4, 12) |  |
| 11 | Gain/Loss | Male and Female | Male | 31 (11, 20) |  |
| 12 | Gain/Loss | Male and Female | Female | 35 (8, 27) |  |
| 13 | Gain/Loss | Male | Unbiased | 1049 (188, 861) | } $p < 0.0001$ |
| 14 | Gain/Loss | Female | Unbiased | 872 (161, 711) |  |
| 15 | Gain/Loss | Male and Female | Unbiased | 53 (12, 41) |  |
| 16 | Gain/Loss | Unbiased | Male | 657 (108, 549) | } $p \approx 0.0001$ |
| 17 | Gain/Loss | Unbiased | Female | 525 (84, 441) |  |
| 18 | Gain/Loss | Unbiased | Male and Female | 27 (5, 22) |  |
| 19 | Expressed | 11716 (1919, 9797) | 11543 (1893, 9650) | 9508 (1529, 7979) |  |

**Supplementary Table 2.**
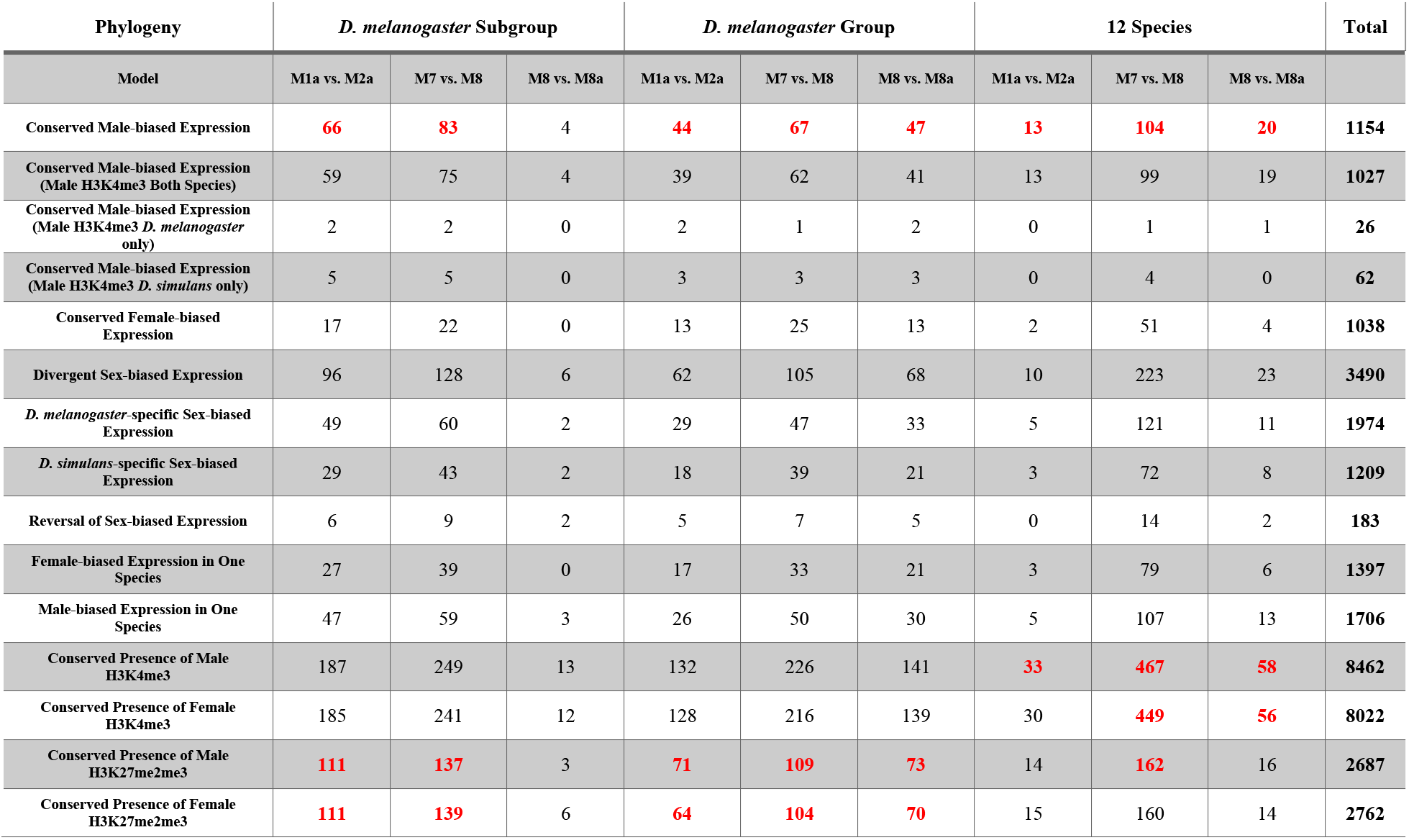
Enrichment of genes with positive selection. Summary of enrichment tests performed between genes with evidence of positive selection from flyDIVas (Stanley and Kulathinal 2016; Clark 2007) and genes with conserved/diverged expression or conserved presence of chromatin marks described in this study. The number of genes with evidence of positive selection for the 3 phylogenetic levels (*D. melanogaster* subgroup, *D. melanogaster* group, and 12 species) and 3 models tested (M1a vs. M2a, M7 vs. M8, and M8 vs. M8a) in flyDIVas is provided for each group. Gene numbers in red are those that were significantly enriched (χ^2^: *p* < 0.001) for genes with positive selection. More detailed descriptions of the models tested can be found in Table 2 of the PAML manual (http://abacus.gene.ucl.ac.uk/software/pamlDOC.pdf). Briefly, M1a vs. M2a compares nearly neutral evolution and positive selection, M7 vs. M8 compares where dN/dS (ω) varies according to a beta distribution vs. a beta distribution plus a discrete ω class where ω > 1 (positive selection), and M8 vs. M8a which compares where ω varies according to a beta distribution with a discrete ω class where ω > 1 vs. a beta distribution with ω=1.

**Supplementary Table 3.**
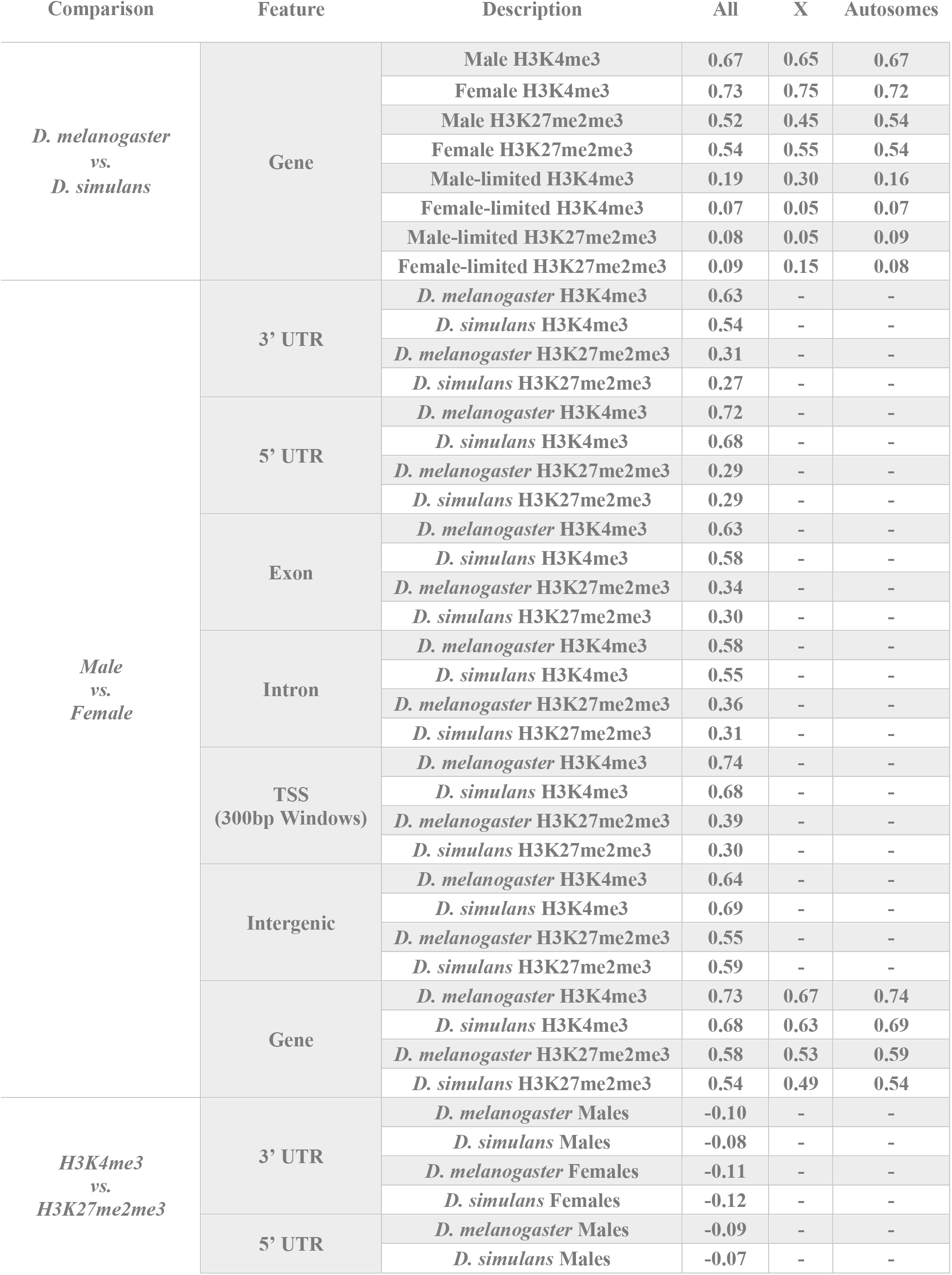
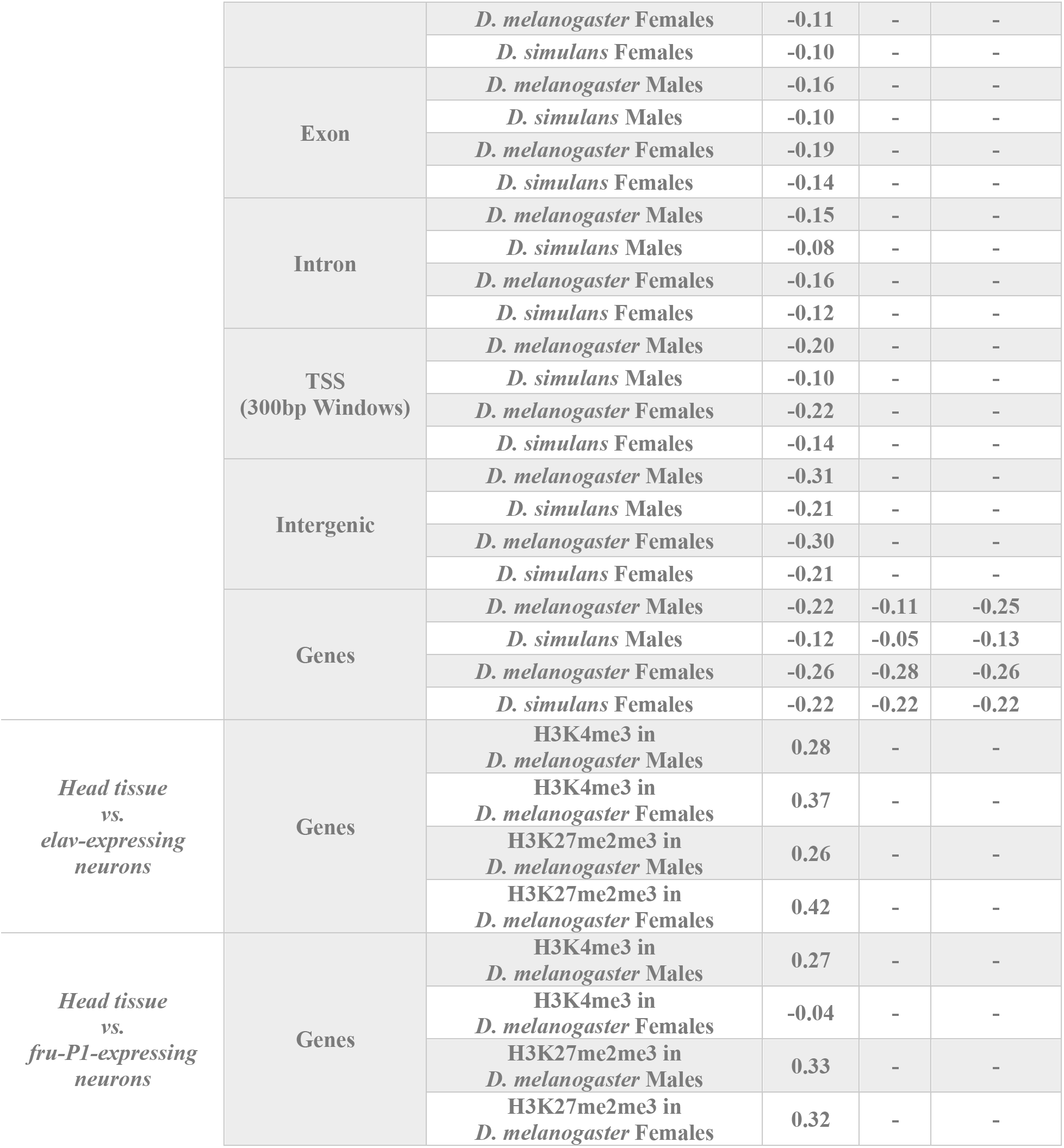
Summary of Kappa values for the indicated comparisons. Cohen’s Kappa values (Fleiss 1981) indicating chance corrected agreement of the comparison described the “Comparison” column for the feature described in the “Feature” column and group in the “Description” column. Kappa values are presented for all chromosomes (X and autosomes combined) for all comparisons, as well as X chromosomes and autosomes separately for several indicated comparisons. Chromosome 4 is excluded from the autosomes.

**Supplementary Table 4.**
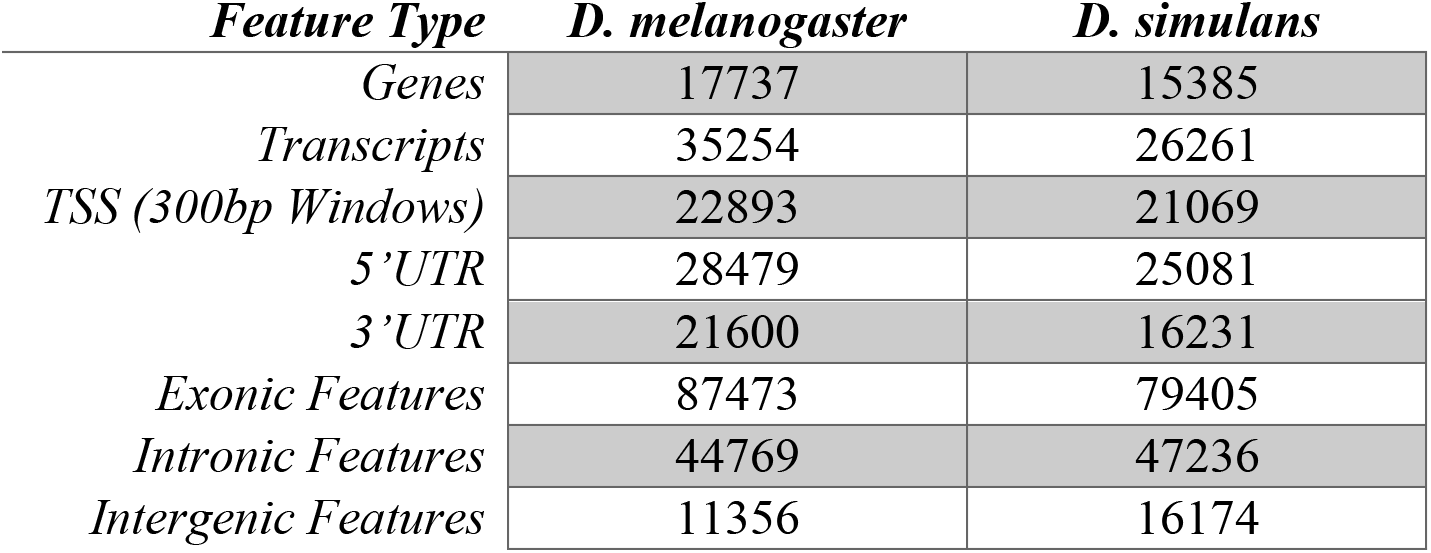
Number of annotated genomic features in *D. melanogaster* and *D. simulans*. Counts of features within *D. melanogaster* and *D. simulans* genome annotation files. 5’ UTR and 3’UTR were determined for each transcript using the references described in the Genome Annotation section of the Methods. A transcription start site (TSS) was defined as a 300 bp region, 150 bp upstream and downstream from each annotated transcript start. In *D. melanogaster* there were three pairs of genes where the members in each pair had the same start position but opposite strands: i) *bug* (FBgn0034050) and *Diap2* (FBgn0015247), ii) *lncRNA:CR44456* (FBgn0265649) and *lncRNA:CR44455* (FBgn0265648), and iii) *CR43482* (FBgn0263493) and *CR43483* (FBgn0263494). Event analysis (Newman, et al. 2018) was used to determine exonic and intronic features. Intergenic features were defined by subtracting the genic features from the entire genome with a length greater than 50 bp.

**Supplementary Table 5.** Summary of features detected by RNA-seq. The number (percent) of features detected in males (irrespective of females), in females (irrespective of males), in either males or females (union), and in both males and females (intersection) for each species mapped to the associated reference genome. Percent (in parentheses) is calculated by dividing the number detected by the total number for each feature type (see Supplementary Table 4 for feature totals).

| Species | Feature Type | # Detected in Males | # Detected in Females | # Detected in Either Sex | # Detected in Both Sexes |
| --- | --- | --- | --- | --- | --- |
| *D. melanogaster* | 3UTR | 14505 (67.15%) | 14280 (66.11%) | 14700 (68.06%) | 14085 (65.21%) |
|  | 5UTR | 18640 (65.45%) | 18198 (63.9%) | 19066 (66.95%) | 17772 (62.4%) |
|  | TSS | 15761 (68.85%) | 15323 (66.93%) | 16161 (70.59%) | 14923 (65.19%) |
|  | Exonic | 69373 (79.31%) | 68038 (77.78%) | 70568 (80.67%) | 66843 (76.42%) |
|  | Intergenic | 6000 (52.84%) | 5633 (49.6%) | 6260 (55.13%) | 5373 (47.31%) |
|  | Intronic | 29555 (66.02%) | 28483 (63.62%) | 30576 (68.3%) | 27462 (61.34%) |
| *D. simulans* | 3UTR | 11820 (72.82%) | 11717 (72.19%) | 12032 (74.13%) | 11505 (70.88%) |
|  | 5UTR | 16108 (64.22%) | 16054 (64.01%) | 16653 (66.4%) | 15509 (61.84%) |
|  | TSS | 13806 (65.53%) | 13712 (65.08%) | 14291 (67.83%) | 13227 (62.78%) |
|  | Exonic | 61777 (77.8%) | 61482 (77.43%) | 63283 (79.7%) | 59976 (75.53%) |
|  | Intergenic | 6769 (41.85%) | 6616 (40.91%) | 7144 (44.17%) | 6241 (38.59%) |
|  | Intronic | 30010 (63.53%) | 30013 (63.54%) | 31531 (66.75%) | 28492 (60.32%) |

**Supplementary Table 6.** Summary of read mapping counts and percentages. (A) RNA-seq mapped reads. All RNA-seq samples were mapped to both the *D. melanogaster* FlyBase 6.17 genome and the *D. simulans* FlyBase r.202 genome. The mean number and percent of processed reads across replicates after mapping to the indicated genome are given. (B) ChIP-seq mapped reads. All ChIP-seq samples were mapped to the associated reference genome based on the species of the sample (*D. melanogaster* FlyBase 6.17 genome or *D. simulans* FlyBase r.202). The mean number and percent of mapped processed reads across replicates for the indicated ChIP mark or input control are given.

| A. |  |  |  |  |  |  |
| --- | --- | --- | --- | --- | --- | --- |
| Species | *D. melanogaster* |  |  |  |  |  |
| Genome | *D. melanogaster* FB r6.17 |  | *D. simulans* FB r2.02 |  |  |  |
| Sex | Male | Female | Male | Female |  |  |
| Mean # mapped reads per replicate | 16,368,252 | 16,479,280 | 16,877,562 | 16,915,308 |  |  |
| Mean % mapped reads per replicate | 91.37% | 92.76% | 94.35% | 95.22% |  |  |
| Species | *D. simulans* |  |  |  |  |  |
| Genome | *D. melanogaster* FB r6.17 |  | *D. simulans* FB r2.02 |  |  |  |
| Sex | Male | Female | Male | Female |  |  |
| Mean # mapped reads per replicate | 14,774,793 | 15,598,854 | 15,498,853 | 16,423,578 |  |  |
| Mean % mapped reads per replicate | 90.44% | 94.61% | 94.83% | 94.61% |  |  |
| B. |  |  |  |  |  |  |
| Species | *D. melanogaster* |  |  |  |  |  |
| Sex | Male |  |  | Female |  |  |
| ChIP/Input | Input | H3K4me3 | H3K27me2me3 | Input | H3K4me3 | H3K27me2me3 |
| Mean # mapped reads per replicate | 10,915,497 | 13,688,173 | 12,666,807 | 12,239,391 | 13,817,377 | 14,380,575 |
| Mean % mapped reads per replicate | 78.82% | 86.08% | 76.38% | 81.83% | 87.73% | 78.70% |
| Species | *D. simulans* |  |  |  |  |  |
| Sex | Male |  |  | Female |  |  |
| ChIP/Input | Input | H3K4me3 | Input | H3K4me3 | Input | H3K4me3 |
| Mean # mapped reads per replicate | 10,435,104 | 14,728,518 | 10,435,104 | 14,728,518 | 10,435,104 | 14,728,518 |
| Mean % mapped read per replicate | 80.25% | 92.01% | 80.25% | 92.01% | 80.25% | 92.01% |

## Supplementary Files

**Supplementary File 1** - Gene-level expression and chromatin accessibility results for *D. melanogaster*. All column variables are defined in Supplementary File 9.

**Supplementary File 2** – Gene-level expression and chromatin accessibility results for *D. simulans*. All column variables are defined in Supplementary File 9.

**Supplementary File 3** – ChIP-seq protocol

**Supplementary File 4** – Orthologous gene pairs of *D. melanogaster* to *D. simulans* selected from FlyBase OrthoDB report (Waterhouse, et al. 2013) in release 2017_04 (dmel_orthologs_in_drosophila_species_fb_2017_04.tsv.gz, downloaded 4/17/19). The original FlyBase file was modified to have individual columns for coordinates, +/- values for strand (compared to 1/-1), and “Dsim\” removed from Ortholog_GeneSymbol elements.

**Supplementary File 5** – Upper quartile values used in for RNA-seq quantification.

**Supplementary File 6** – Feature-level expression and chromatin accessibility results for *D. melanogaster*.

**Supplementary File 7** – Feature-level expression and chromatin accessibility results for *D. simulans*.

**Supplementary File 8** – Gene-level expression and chromatin accessibility results for *D. melanogaster* and *D. simulans* orthologs as identified by the FlyBase OrthoDB report (Waterhouse, et al. 2013). All column variables are defined in Supplementary File 10.

**Supplementary File 9** – Gene-level variable definitions for species result files (Supplementary Files 1, 2).

**Supplementary File 10** – Gene-level variable definitions for the *D. melanogaster* and *D. simulans* ortholog result file (Supplementary File 8).

**Supplementary File 11** – For all gene numbers called out in the main text, the descriptions and the flags needed to identify those genes in Supplementary Files 1 or 8.

